# A kinetic dichotomy between mitochondrial and nuclear gene expression drives OXPHOS biogenesis

**DOI:** 10.1101/2023.02.09.527880

**Authors:** Erik McShane, Mary Couvillion, Robert Ietswaart, Gyan Prakash, Brendan M. Smalec, Iliana Soto, Autum R. Baxter-Koenigs, Karine Choquet, L. Stirling Churchman

**Author notes:** Corresponding author: L. Stirling Churchman Department of Genetics, Blavatnik Institute, Harvard Medical School, Boston, MA 02115.

## Abstract

Oxidative phosphorylation (OXPHOS) complexes, encoded by both mitochondrial and nuclear DNA, are essential producers of cellular ATP, but how nuclear and mitochondrial gene expression steps are coordinated to achieve balanced OXPHOS biogenesis remains unresolved. Here, we present a parallel quantitative analysis of the human nuclear and mitochondrial messenger RNA (mt-mRNA) life cycles, including transcript production, processing, ribosome association, and degradation. The kinetic rates of nearly every stage of gene expression differed starkly across compartments. Compared to nuclear mRNAs, mt-mRNAs were produced 700-fold higher, degraded 5-fold faster, and accumulated to 170-fold higher levels. Quantitative modeling and depletion of mitochondrial factors, LRPPRC and FASTKD5, identified critical points of mitochondrial regulatory control, revealing that the mitonuclear expression disparities intrinsically arise from the highly polycistronic nature of human mitochondrial pre-mRNA. We propose that resolving these differences requires a100-fold slower mitochondrial translation rate, illuminating the mitoribosome as a nexus of mitonuclear co-regulation.

## Introduction

Eukaryotic cells maintain and express two genomes: the nuclear chromosomes and the small, circular, and highly polyploid mitochondrial DNA (mtDNA). Mitochondrial and nuclear gene expression processes are spatially separated and governed by evolutionary distinct transcription, RNA processing, and translation machinery^1^. The human mitochondrial genome encodes subunits of the oxidative phosphorylation (OXPHOS) complexes responsible for cellular respiration and ATP production^1–3^. As the nuclear genome encodes the rest of the OXPHOS subunits, nuclear and mitochondrial DNA expression must be coordinated, as evidenced by observations of balanced human OXPHOS protein synthesis across compartments^3^. Dysregulation of mitonuclear protein synthesis causes proteostatic stress^3,4^, and mutations in genes involved in mitochondrial gene expression can cause severe human disease syndromes^5^.

Nuclear-encoded genes are transcribed from two DNA copies into monocistronic transcripts that are spliced, polyadenylated, and exported into the cytoplasm for translation. By contrast, mitochondrial genes are transcribed from hundreds of copies of 16.5 kb mtDNA into two long polycistronic primary transcripts encoded on each strand, the heavy and light strands (Fig 1A)^6^. The primary transcripts are cleaved into individual mt-mRNAs, transfer RNAs (mt-tRNA), and ribosomal RNAs (mt-rRNA)^7,8^. After processing, mt-mRNA is typically adorned with short poly(A) tails before being translated by the mitochondrial ribosomes (mitoribosomes) at the inner mitochondrial membrane^9^. Because mt-RNAs are produced from only three promoters^10^, regulation of transcription initiation is presumed to contribute little to the relative levels of mitochondrial-encoded proteins. Rather, the steady-state mitochondrial transcriptome is thought to be shaped by post-transcriptional processes. This gene expression strategy is distinctive to bilaterian mitochondria^11,12^. For example, *S. cerevisiae* mtDNA has 11 transcription start sites such that rRNA production is largely uncoupled from OXPHOS mRNA production^13^. Furthermore, despite their alphaproteobacterial origin, human mitochondria use polycistronic transcription differently from *E. coli,* where individual mRNAs are not cleaved out of primary transcripts^14^. Thus, the evolutionary outcome for animal mitochondria results in a gene expression system with distinctive challenges due to the intrinsic coupling of coding and non-coding RNA production. These differences suggest that the life cycles of mitochondrial RNA must be dramatically different to those of nuclear RNA. However, mt-RNAs biogenesis, accumulation and ribosome association have not been described as quantitatively as for nuclear RNAs, hindering a precise comparison of OXPHOS gene expression across compartments. To understand OXPHOS biogenesis, we need to quantitatively compare the nuclear and mitochondrial gene expression systems to determine how they resolve their differences to coordinate the synthesis of OXPHOS subunits.

**Figure 1.**
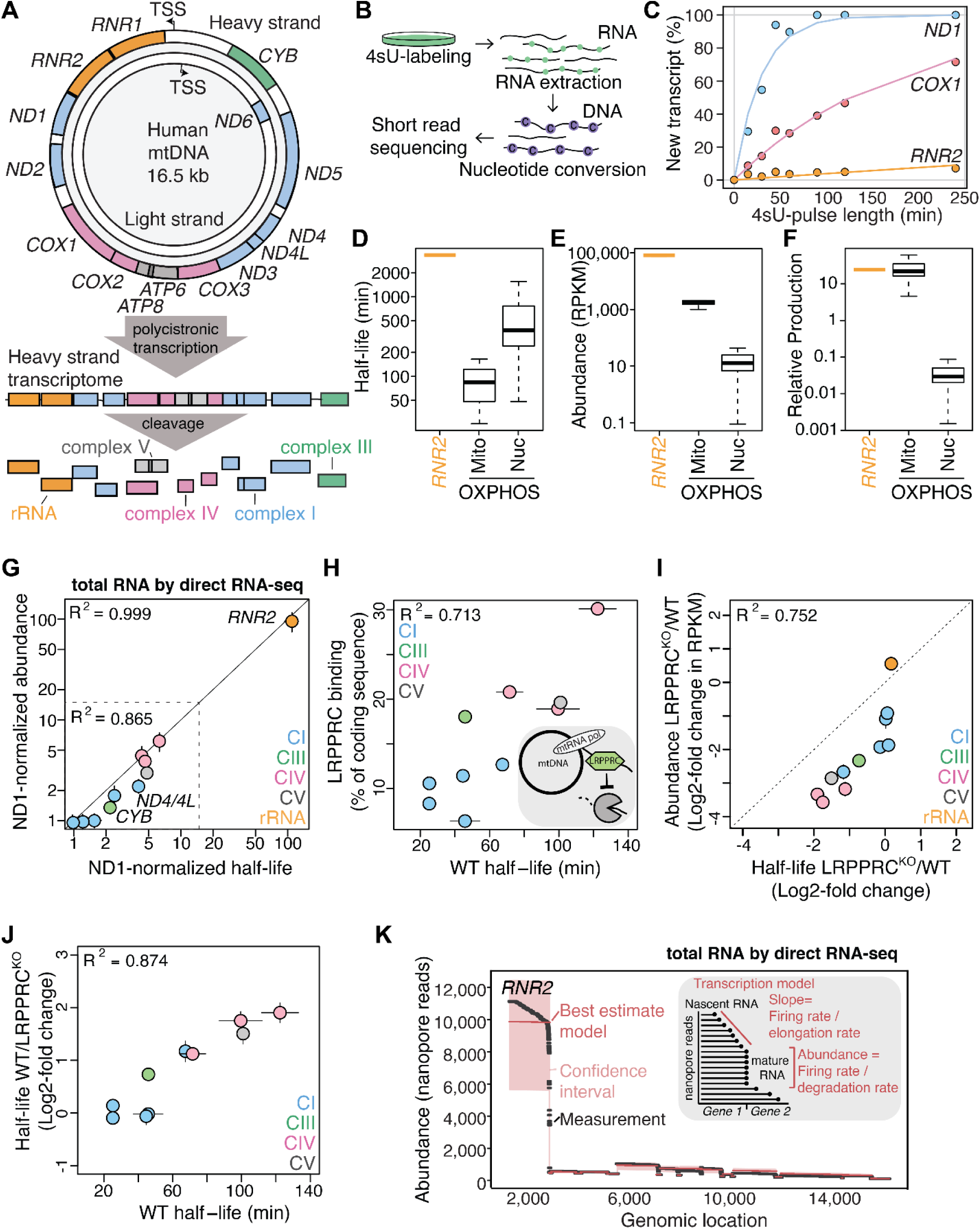
RNA turnover is set by LRRPRC and is the main determinant of the relative abundance of the heavy strand–encoded transcriptome. A) The heavy strand of the mitochondrial genome encodes two ribosomal RNAs, 10 mt-mRNAs (two bicistronic), and several tRNAs, whereas the light strand encodes one mRNA as well as tRNAs. Mitochondrial transcription is polycistronic, with all transcription start sites located upstream of the coding regions^10^, resulting in long primary transcripts that are cleaved by processing enzymes into individual rRNAs, tRNAs, and mRNAs (color-coded by OXPHOS complex). B) Overview of TimeLapse-seq, which produces measurements of fraction new RNA. Green dots indicate metabolically incorporated 4sU, which is modified during TL chemistry and misread as C (purple dots) during reverse transcription. C) Turnover profiles for two mt-mRNAs, *COX1* and *ND1*, and one mt-rRNA, *RNR2*. Dots show the measured data, and lines indicate the best fit for an exponential model. D) Distribution of half-lives derived from the exponential model for the mt-rRNA *RNR2*, all heavy strand–encoded mt-mRNAs (“Mito”, n=10), and all nuclear mRNAs encoding OXPHOS subunits (“Nuc”, n= 76). E) Same as D), but for transcript abundance estimated based on the RPKMs from the same experiment. Zero values were excluded. F) Production rates calculated from the turnover rates in D) and the abundances in E) by dividing the RPKM by the half-life. G) *ND1*-normalized abundance from nanopore direct RNA-seq compared to *ND1*-normalized turnover rates from mito-TL-seq. Error bars represent standard deviation. Pearson R^2^, calculated in non-log space, is displayed for all transcripts (top left corner) and all mt-mRNAs (excluding *RNR2*, top left corner in dashed line box). Two promoter distal genes are highlighted. H) RNAse footprinting data from mouse embryonic fibroblasts (MEF)^27^ showing the fraction of each mRNA that was bound by LRPPRC. LRPPRC binding was compared to the half-lives derived from mito-TL-seq. I) Scatter plot showing the correlation between the change in RNA abundance and the change in turnover in HEK293T cells depleted of LRPPRC (LRPPRC^KO^) relative to wild-type (WT) cells. Data from two replicates are shown. Error bars show the standard deviation. The Pearson R^2^ is shown. J) Scatter plot showing the positive correlation between mRNA half-lives in HEK293T wild-type cells and change in stability in cells containing LRPPRC relative to knockout cells. The most stable mRNAs are most affected by the loss of LRPPRC. The Pearson R^2^ is shown. K) Reads from one direct RNA-seq experiment mapping to the mitochondrial heavy strand (in black), with the results from the transcription model (shown in the right red box) overlaid in red. The model takes into account turnover and transcription parameters to estimate RNA abundance. Firing rate = slope of 3’ ends of mapped reads from direct RNA-seq/elongation rate^28,29^. All data in the figure are from HeLa cells except H) which is from MEF, and I) and J) which are from HEK293T cells.

Here, we developed numerous approaches tailored to the mitochondrial genome to quantitatively determine each stage of the mt-mRNA life cycle. Through comparing nuclear-and mitochondrial-encoded mRNAs, we found a striking discordance at all major stages of gene expression across compartments. In contrast to nuclear-encoded genes, differential turnover explains steady-state mt-mRNA abundance and is set largely by LRPPRC, a protein associated with the metabolic disorder Leigh syndrome^15^. Ribosome association occurred 20-fold faster for mt-mRNA and 5’-end mt-mRNA processing out of the polycistronic transcript was nearly obligatory for mitoribosome association, even when processing is severely disrupted upon loss of FASTKD5. We combined our kinetic measurements to develop a quantitative gene expression model and showed that mitochondrial and nuclear protein synthesis experience the most regulation through RNA turnover and transcription initiation, respectively, with an additional contribution from translation regulation for both.

With these wide differences across the gene expression systems, a resolution is required to balance OXPHOS subunit synthesis. We propose that this balance comes from slow mitochondrial translation rates, orders of magnitude slower than cytosolic translation. Our model of mitochondrial and nuclear gene expression control is the first systematic, quantitative, and comparative kinetic analysis of the dual-origin OXPHOS transcriptomes. By revealing the necessary points of mitochondrial and nuclear co-regulation, these results shed light on the unique vulnerabilities of human mitonuclear balance that may contribute to disease and aging processes.

## Results

### Mitochondrial mRNA degradation rates are 5-fold faster than nuclear-encoded genes

The mtDNA heavy strand encodes two mitochondrial rRNAs, *RNR1* and *RNR2*, and 10 of the 11 mt-mRNAs (Figure 1A), which are produced in a single primary transcript and processed into individual RNAs^7^. Heavy-strand RNAs accumulate at levels that vary over two orders of magnitude despite being produced in equimolar proportions^16,17^. RNA stability has been proposed to be a leading contributor to the differential steady-state levels of the individual transcripts^18,19^. However, testing this hypothesis has been challenging due to a lack of methods to quantify RNA turnover that does not rely on strong perturbations, e.g., transcription inhibition, which leads to compensatory changes to RNA turnover rates^16,18^. Recent advances in the detection of modified bases by next-generation sequencing have led to the development of nucleotide-conversion-based strategies to measure RNA turnover (Figure 1B)^20–22^. We leveraged this strategy and adapted TimeLapse-seq (TL-seq)^21^ for the quantitative measurement of mitochondrial RNA (mito-TL-seq). In our approach, the nucleotide analog 4-thiouridine (4sU) is introduced to cells for various amounts of time. After RNA extraction, incorporated 4sU is chemically modified, causing it to be misread during reverse transcription and resulting in T-to-C mismatches during sequencing (Figure 1B). The fraction of new transcripts for each gene is determined by the analysis of mismatches that occur after sequence alignment, and a simple exponential decay model is used to estimate the half-lives of all transcripts (Figure 1C).

In developing mito-TL-seq, we found that 4sU was incorporated at a lower frequency in mitochondrial-encoded RNAs than in nuclear-encoded transcripts (Figure S1A). Because high levels of 4sU lead to numerous secondary effects^23^, rather than increase the concentration of 4sU, we re-established computational approaches used in analyzing these data^24,25^ to properly quantify T-to-C conversions in mitochondrial RNA sequencing reads. We developed strategies to decrease background, including creating reference genomes where single nucleotide polymorphisms have been removed, analyzed reads aligning to the mitochondrial genome separately, and determined the T-to-C conversion rates using a custom binomial mixture model^25^. Using this approach, we analyzed HeLa cells labeled with a 4sU concentration that causes minimal secondary effects (50 uM)^23^. Turnover rates were reproducible between replicates and independent of 4sU concentration (Figures 1C-D, S1D, and Table S1). Moreover, we confirmed these results with an alternative method that did not rely on detecting 4sU-RNA or sequencing which used the probe-based approach, MitoStrings (Figures S1B-D, and Table S1)^25,26^. We observed a strong correlation between the MitoStrings and mito-TL-seq half-lives (Figure S1D, R^2^ > 0.9).

We compared the half-lives of nuclear-and mitochondrial heavy strand–encoded OXPHOS mRNA, as well as *RNR2* (*RNR1* is removed during an rRNA depletion step). Half-lives ranged broadly, with mitochondrial-encoded mRNA (median half-life of 379 min) turned over about 5-fold more rapidly than nuclear-encoded OXPHOS mRNAs (median half-life of 84 min) (Figures 1D, and S1R). By contrast, *RNR2* was turned over >20-fold slower than any OXPHOS mRNA. At steady-state, mt-mRNAs were ∼170-fold more abundant than nuclear-encoded OXPHOS mRNAs (Figures 1E, and S1R, and Table S2) and *RNR2* was 50-fold more abundant than the mt-mRNAs (Figure 1E). To estimate production levels for each RNA, we divided the abundance estimates by the half-lives for each gene (Figures 1F, and S1R). Importantly, production rates for mitochondrial rRNA and mRNAs were comparable (Figures 1F, and S1R), consistent with the polycistronic nature of mitochondrial transcription.

However, this coupling of coding and non-coding RNA led to mitochondrial mRNA production rates 700-fold higher than for nuclear-encoded OXPHOS mRNAs.

### LRPPRC-controlled RNA turnover rates set mitochondrial mRNA levels

We sought to determine to what extent mt-mRNA stability was responsible for the differential RNA abundances across the mitochondrial transcriptome. We used nanopore direct RNA sequencing to obtain high accuracy measurements of mt-RNA abundance with minimal biases from library-generation steps, such as PCR amplification (Figures 1G, and S1E, Table S2). By comparing the half-lives from mito-TL-seq with the abundance from direct RNA-seq, we found that the coefficient of determination (R^2^) is 0.858, indicating that more than 85% of the variability in mRNA abundance can be explained by degradation alone (Figure 1G). Notably, the dynamic ranges of each measurement are matched (with *ND1*-relative abundance and half-lives both ranging between 0 to 100), which must be the case if turnover is truly a driver of the steady-state abundance of mitochondrial RNAs. We obtained similar results for the light-strand transcriptome, despite technical limitations in quantifying light-strand encoded RNA abundances (see Methods, and Figure S1F-N).

As turnover helps establish the mt-mRNA steady-state levels, we next wondered what is responsible for the heterogeneity in mt-mRNA half-lives. Mutations in *LRPPRC* cause the French Canadian type of Leigh syndrome, and the LRPPRC protein is involved in mt-mRNA stabilization, poly(A) tailing, and translation in both mice and humans (Figure 1IH)^3,15,30,31^. Intriguingly, we found a strong correlation between the fraction of each mRNA bound by LRPPRC^27^ and their turnover rates (R^2^ = 0.713) (Figure 1H), indicating that LRPPRC may set RNA degradation rates. To test this hypothesis, we analyzed *LRPPRC* knockout and matched control cells^3^. Depletion of *LRPPRC* had profound effects on the steady-state levels of most heavy-strand encoded RNAs (Figure 1I). Emphasizing the role of RNA turnover in setting steady-state abundances of mt-mRNAs, we found that changes in RNA half-lives after *LRPPRC* depletion correlated with the changes in the steady-state levels of RNA (R^2^ = 0.752) (Figure 1I). To determine the role of LRPPRC in establishing mt-mRNA stability, we compared the stabilities of wild-type mt-mRNAs with the stability enhancement due to the presence of LRPPRC and observed a strong positive correlation, indicating that LRPPRC sets 87% of steady-state half-life variability (Figure 1J). Taken together, we conclude that differential mt-mRNA abundances are explained largely by LRPPRC-dependent RNA turnover.

### Slow mt-RNA transcription elongation contributes to RNA abundances

Our measurements of differential RNA turnover did not completely explain the relative abundances of mitochondrial RNA. The transcripts that deviated from a direct one-to-one correspondence between turnover and abundance levels were often encoded on the promoter-distal end (e.g. *CYB* and *ND4/4L*) (Figure 1G). When transcription is polycistronic, more actively transcribing RNA polymerases will have traversed promoter-proximal genes than promoter-distal genes at any point in time (Figure S1O). We reasoned that if transcription is slow but RNA turnover is fast, transcription elongation will lead to more promoter-proximal than distal nascent RNA. We used mathematical modeling^28^ to predict total RNA abundance from the estimates of mature RNA levels using our turnover measurements and the nascent RNA levels (Figures 1K, and S1P). The model sucessfully explained the remaining differences in mt-mRNA abundances and placed an mt-RNA polymerase elongation rate in the range 0.06–0.12 kb/min (Figures 1K, and S1P,Q), on order with *in vitro* mitochondrial transcription rate estimates (0.22 to 0.72 kb/min^32–34)^. In summary, our kinetic measurements suggest that the steady-state mitochondrial transcriptome is shaped primarily by differential degradation, with transcription elongation providing a differential contribution for genes encoded proximal to the heavy strand transcription start site.

### Mitochondrial RNA processing is rapid and occurs predominantly co-transcriptionally

Most nuclear pre-mRNA splicing events occur co-transcriptionally^35,36^, with most introns removed several minutes or kilobases after their synthesis^37,38^. Accordingly, introns in nuclear-encoded OXPHOS mRNA take around 9 minutes to be excised^37^, and the mRNA remains on chromatin for an average of 35 minutes (Figure 6A)^25^. Mitochondrial precursor RNAs are processed into individual RNA molecules mainly through the excision of tRNAs that are encoded at the 5’ and 3’ ends of mRNAs^7^. Considering that mt-mRNA are turned over rapidly, processing must occur faster than for nuclear genes or alternatively, be dispensable for mt-mRNA function. Therefore, we next investigated how mt-mRNA processing kinetics relate to nuclear processing rates and mitochondrial transcription and turnover rates. To calculate processing rates, we used nanopore direct RNA sequencing reads to quantitatively capture both processed and unprocessed transcripts (Figures 2A-B, and S2A). These data enabled measurements of the fraction of unprocessed transcripts at steady state, which we used to estimate processing rates (Figure 2C, and Table S3) and confirmed for some transcripts by MitoStrings (Figure S2F, Material and Methods). Our analysis revealed that processing generally occurs rapidly, with most processing half-lives being less than 1 minute (Figures 2D, and S2B). Some processing events took considerably longer (9–39 min, Figure 2D). In one expected case, the processing of *ATP8/6* and *COX3*, which was previously observed to accumulate as a translation-competent tricistronic transcript^3,26,39^, took over 20 minutes (Figure 2A, D). Taken together, because the mitochondrial transcription elongation rate is less than 1 kb/min (Figure 1I), we conclude that mitochondrial processing must occur predominantly co-transcriptionally, similar to nuclear pre-mRNA splicing^35,36^.

**Figure 2.**
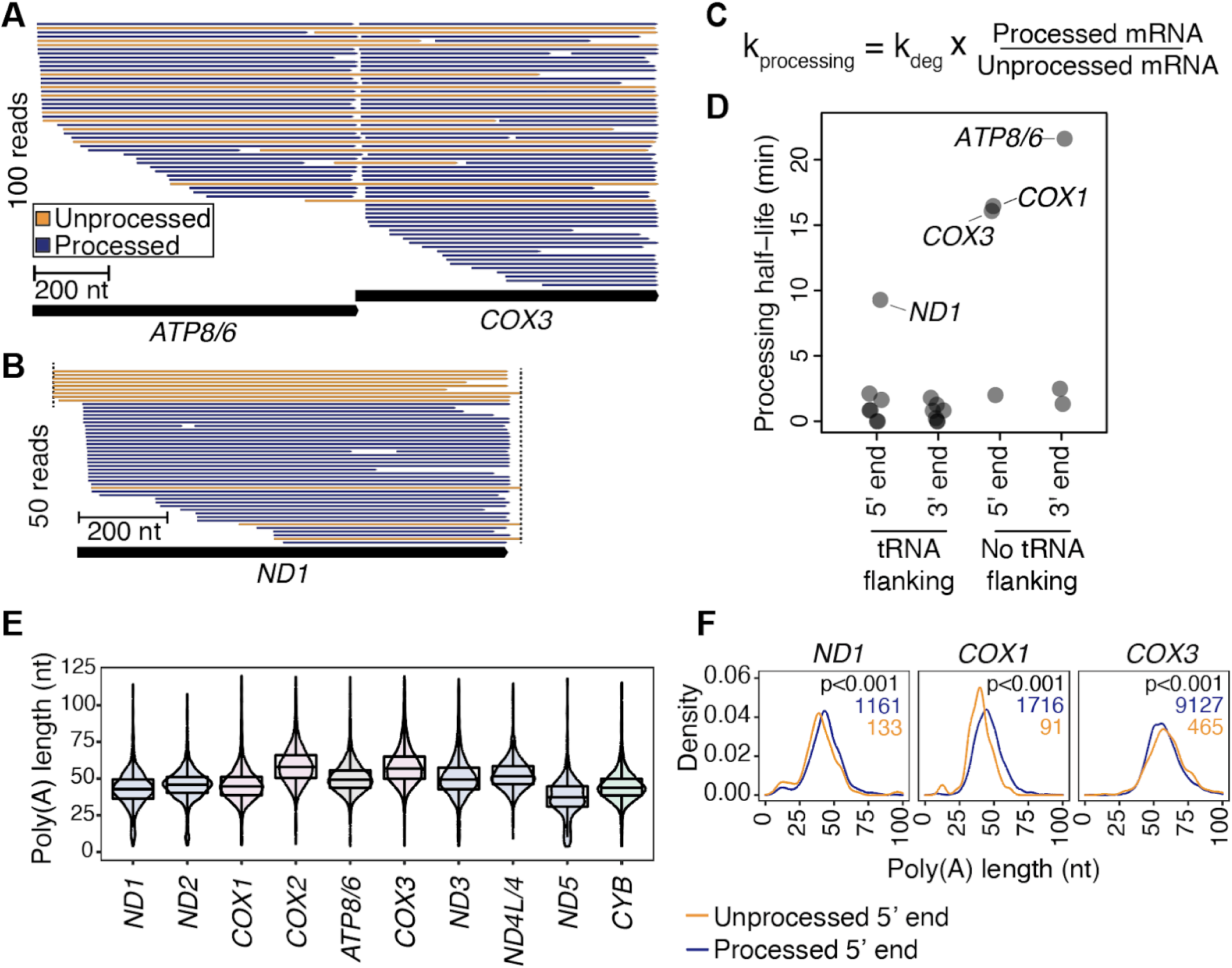
Mitochondrial RNA processing is generally rapid, with notable exceptions. A) 100 randomly sampled reads from nanopore direct RNA-seq that align to *ATP8/6* and/or *COX3* include many transcripts (orange reads) that are unprocessed at the 5’ and/or 3’ ends. B) Same as A), but for 50 reads mapping to *ND1*. C) Equation used to estimate processing rates from nanopore direct RNA seq and mito-TL-seq turnover measurements as shown in D). D) The amount of time required until half of the newly synthesized transcripts have been processed at a specific site. Processing sites are subdivided into 3’ or 5’ ends of the mRNA, as well as into groups of canonical (tRNA-flanking) or non-canonical junctions. E) Distribution of poly(A) tail lengths measured by direct RNA-seq of poly(A)+ RNA. All pairwise comparisons between transcripts are significant (p-value < 0.05) by the Wilcoxon rank-sum test, except *ND3* vs. *ATP8/6*. F) Density plot showing the distribution of poly(A) tail lengths for three genes. Orange lines show mRNAs with unprocessed 5’ ends (more nascent on average) and blue lines show mRNAs with processed 5’ ends. The numbers of reads underlying each distribution (in blue or yellow, respectively) and the p-value from the Wilcoxon rank-sum test comparing processed and unprocessed reads are displayed. All data in the figure are from HeLa cells.

### Mitochondrial poly(A) tailing occurs rapidly after 3’-end processing

Mitochondrial mRNAs, except *ND6*, have short, 40-60 nt long, poly(A) tails^40^ compared to nuclear-encoded mRNA (110 nt long)^25,41^. For most mt-transcripts, the poly(A) tail is required to complete the stop codon^40^, but some reports suggest that it is dispensable^42,43^. We wondered if poly(A) tailing kinetics were under less stringent time requirements and would occur more slowly than processing. To test this hypothesis, we used nanopore direct RNA sequencing to quantify the proportion of RNAs with and without a poly(A) tail by using a modified protocol that captured all RNAs regardless of 3’ end status. Most mRNAs had poly(A) tail lengths between 40 and 60 nucleotides, consistent with previous studies (Figures 2E, and S2G, and Table S4). Interestingly, only a small fraction of completed transcripts (< 4%) lacked poly(A) tails, indicating that they are added quickly (Figure S2H). Due to the low level of non-polyadenylated reads, we obtained insufficient coverage to calculate polyadenylation rates. Nevertheless, our results indicated that polyadenylation occurs very rapidly following 3’-end cleavage, similar to nuclear polyadenylation^44^.

To analyze the timing of 3’-end polyadenylation relative to 5’-end processing, we compared the lengths of poly(A) tails as a function of the 5’ processing status. We saw a significant difference for several mRNAs (Figures 2F, and S2I). *ND1*, *COX1,* and *CYB* mRNAs with unprocessed 5’-ends had shorter tails than those with processed 5’-ends, suggesting that 3’ end polyadenylation occurs while 5’ end processing is ongoing. In contrast, *COX3* mRNA showed the opposite trend, with unprocessed transcripts having slightly longer tails (median 2.3 nt longer) than the processed transcripts. If poly(A) tail length reflects the age of the transcripts, this observation is consistent with the higher stability of the tricistronic *ATP8/6-COX3* mRNA compared to the processed *COX3* mRNA. In sum, poly(A) tailing is initiated rapidly on the same time scales as processing^42^.

### mt-mRNAs arrive at ribosomes over 20-fold more rapidly than their nuclear counterparts

The mechanism of mt-mRNA association with the mitoribosome has recently been resolved^45^, however, it remains unknown how long mt-mRNAs take before associating with mitoribosomes and what regulates their arrival time. To address this, we performed mito-TL-seq on RNA associated with mitoribosomes, enriched by sucrose gradient fractionation and immunoprecipitation^3^ (Figures 3A-B, and S3C-D). Due to the challenges of analyzing light-strand RNAs (Figure S1F-N), *ND6* was not included in these analyses.

**Figure 3.**
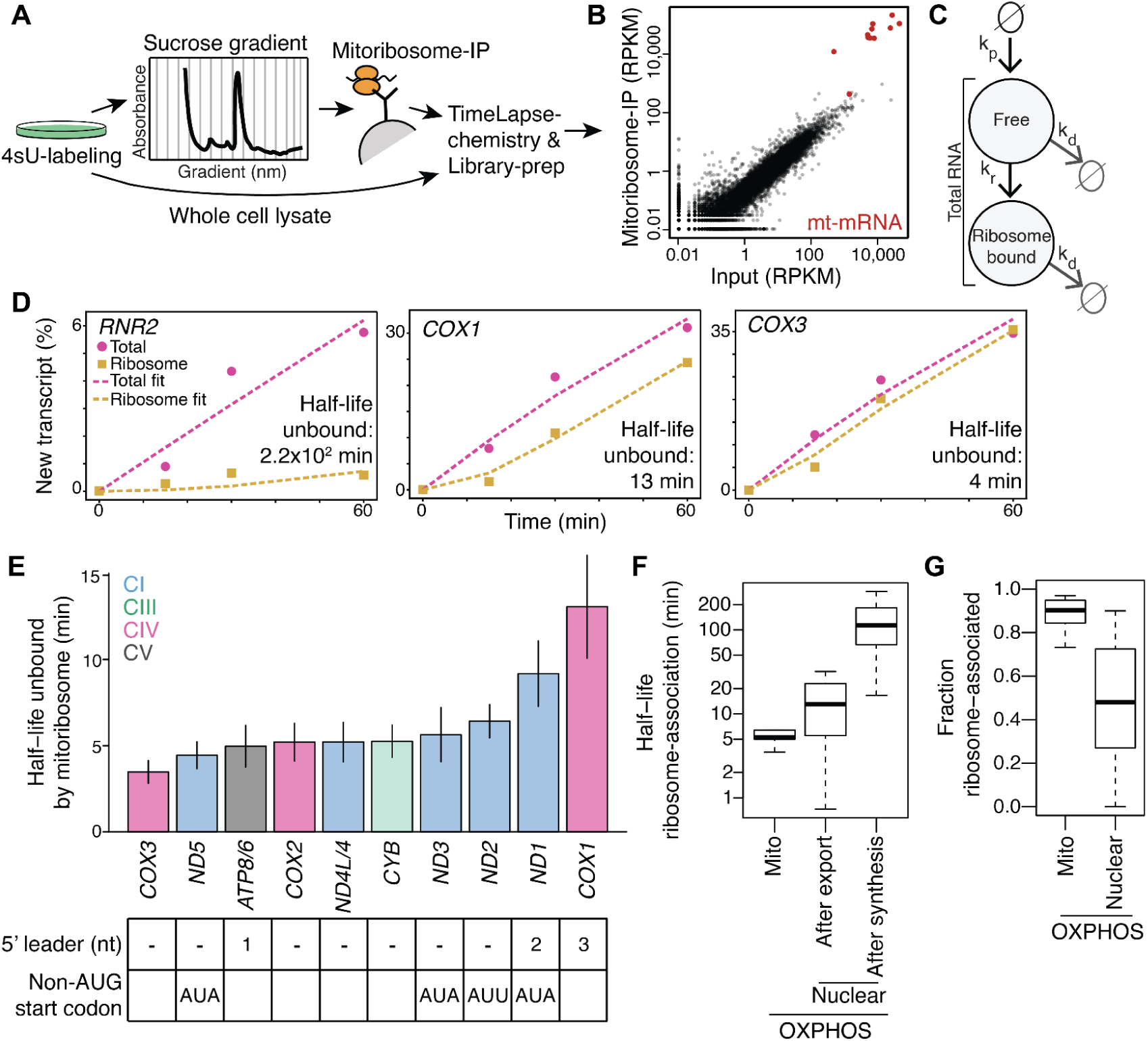
Mitoribosome-association kinetics is significantly faster for mitochondrial-encoded transcripts. A) Cell lysates from 4sU-labeled cells were fractionated using sucrose gradients, and the mitoribosomes were immunoprecipitated (IP) out of the gradient. RNA was extracted from the IPs and input (whole-cell lysate) and sequenced by mito-TL-seq. B) Scatter plot showing the reads per kilobase per million reads (RPKM) in libraries from the mitoribosome-IP experiment and the input samples for the same experiment. Mt-ribosomal mRNA and mt-rRNA are enriched in the IP and shown in red. C) Model used to calculate ribosome association rates. *k*_*deg*_ = degradation rate and *k*_*r*_ = transfer rate from the unbound to the ribosome-bound state. RNA can be either unbound (free) or bound by ribosomes. D) Three example profiles from one replicate experiment performed using 100 µM 4sU. Black squares show the fraction new RNA in the mitoribosome-IP and black circles show the fraction new RNA in the input (Total). Dotted purple and yellow lines show the best fit for the total and mitoribosome-IP, respectively, deploying the model in C). Ribosome-association half-lives (half-life unbound, the time it takes for half of the transcripts to bind to the ribosome), in minutes, are estimated by the model for each gene. E) Barplot showing the time it takes for mt-mRNA to associate with the mitoribosome. mt-mRNA 5’ end characteristics are listed below the plot. Error bars show standard deviations from 3 experiments. F) Distribution of ribosome association half-lives for mitochondrial-encoded mRNA or nuclear-encoded OXPHOS subunits, measured either from nuclear export (shorter half-lives) or from synthesis (longer). Mito data are from this study and HeLa cells, whereas nuclear-encoded data are from^25^ and K562 cells. G) Same as F) but showing the distribution of the fraction of each mRNA that is bound by ribosomes. Mitochondrial mRNAs are on average more frequently associated with the mitoribosome than nuclear-encoded transcripts with the cytosolic ribosomes. All the data in the figure are from HeLa cells if nothing else is stated.

To determine ribosome association rates, we derived a mathematical model (Figure 3C) to describe the flow of mt-mRNAs as they are produced and progress from an unbound (free) state to a ribosome-bound state, assuming equal mRNA degradation rates for both the unbound and ribosome-bound transcripts. An Akaike information criterion (AIC) test consistently preferred this simpler model to a slightly more advanced model with independent degradation rates for the free and ribosome-bound state. Results were very similar regardless of model used (Figure S3E). The mt-rRNA *RNR2* was estimated to take >200 minutes to reach the monosome, consistent with estimates of mitoribosome assembly of 2–3 hours (Figure 3D, and Table S5)^46^. For mt-mRNA, the association times varied, with half-lives of unbound mt-mRNA ranging from 4 minutes for *COX3* to 13 minutes for *COX1* (Figures 3D-E, and S3F, and Table S5). The variability in mitoribosome association rates for mt-mRNA was surprising because no physical barrier, such as the nuclear envelope, separates nascent RNA and ribosomes in mitochondria, and mt-mRNAs have minimal or absent 5’ UTRs. These trends were only weakly related to the size of the 5’ UTR, most of which are 0 or 1 nt, or the start codon (Figure 3E)^47^. Overall, our measurements showed that ribosome association kinetics differed by a factor of >3 across mt-mRNAs.

Due to their compartment-specific transcription, nuclear export, and translation kinetics, nuclear-encoded OXPHOS transcripts spend a long time unbound by ribosomes^25^. Comparing mitochondrial and cytosolic ribosome association rates, we found that nuclear-encoded transcripts spent >20x more time in a non-ribosome-bound state than mitochondrial-encoded transcripts (median time: nuclear-encoded 113 min, mitochondrial-encoded 5 min; Figure 3F). Furthermore, nuclear-encoded mRNAs spent >2x more time between nuclear export and ribosome association compared to the time mt-mRNAs are unbound by ribosomes illustrating how rapid the process is in mitochondria^25^ (median mRNA residence time in the cytoplasm unbound to ribosomes: 13 min, Figure 3F)^25^. Using the ribosome association rates, we calculated the fraction of each mRNA expected to be associated with the ribosome. We found that 90% of mt-mRNA should be ribosome-bound on average, consistent with biochemical measurements^48^ and far higher than for the ∼50% we estimated for nuclear-encoded transcripts (Figure 3G).

### 5’ but not 3’ mt-mRNA processing is a prerequisite for mitoribosome association

Because no physical barriers separate transcription and translation in mitochondria, we sought to determine why ribosome association rates vary. We reasoned that RNA processing, the immediately preceding step, might contribute to this variation, especially as *in vitro* free 5’ ends aid in mitoribosome binding^49^. We found that ribosome association rates (median time until half of mRNA associated with ribosome: 5 min) were generally slower than processing rates (median 5’ and 3’ processing half-life: 100 s), with three exceptions (Figure 4A, B). For both *ND1* and *COX1*, 5’-end processing took the same amount of time as ribosome association, and for the tricistronic *ATP8/6-COX3* transcript, processing was slower than ribosome association (Figure 4A, B). These results led us to hypothesize that for all mt-mRNAs, except for *COX3,* 5’-end processing is a prerequisite of ribosome association *in vivo*.

**Figure 4.**
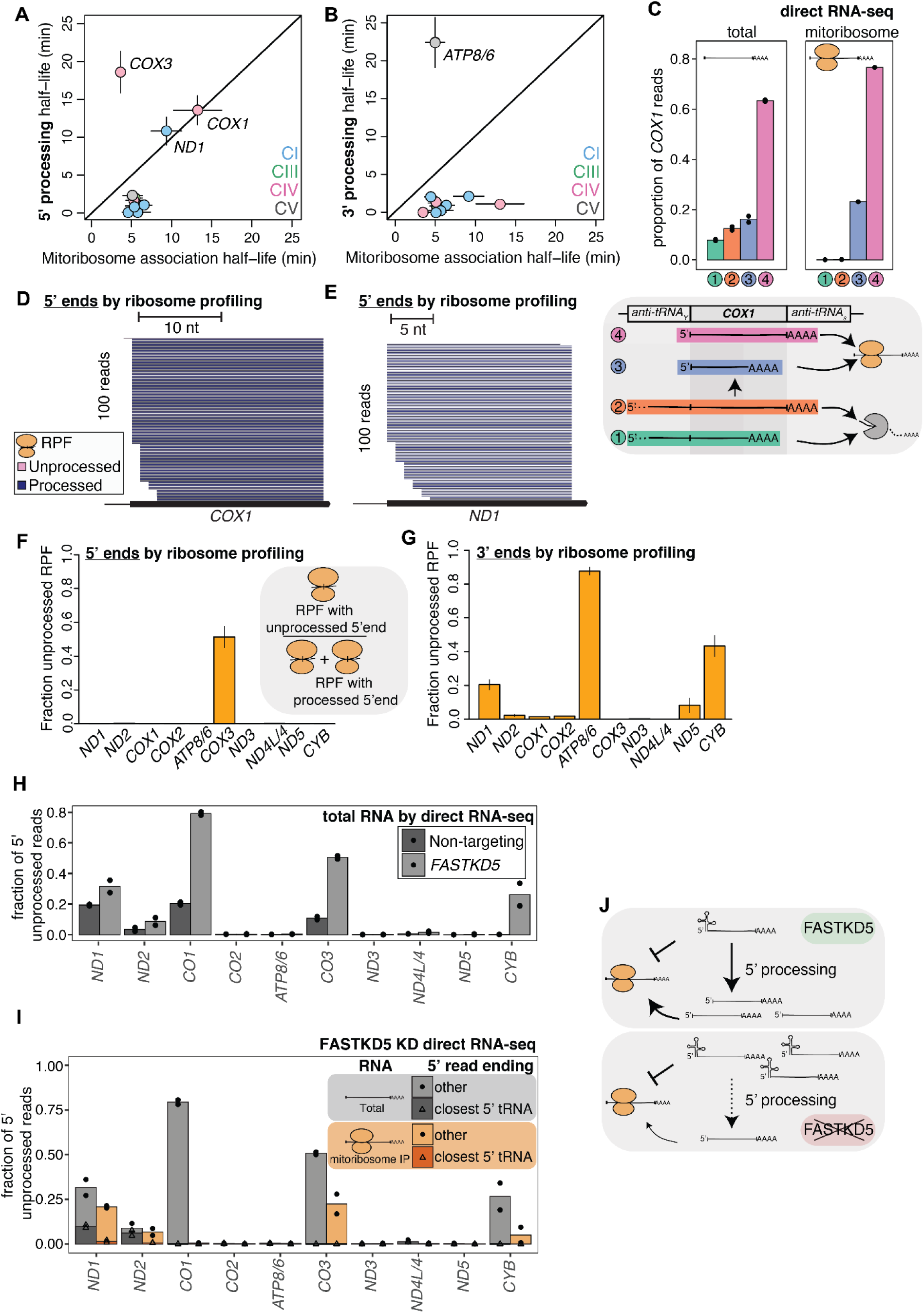
5’ end unprocessed mt-mRNA-processing are depleted from the mitoribosome. A) Scatter plot correlating 5’-end processing and mitoribosome-association half-lives. Error bars show standard deviations for three replicates. B) Same as A) but showing the 3’-end processing half-lives. C) The proportion of *COX1* mRNA based on their 5’-end processing status and 3’-end location as measured by direct RNA-seq of total or ribosome-associated RNA. The scheme below summarizes the finding that only 5’ end processed mRNAs associate with the ribosome. D) Subsampling of 100 ribosome-protected fragments (RPF) with 5’ ends within -30 and +3 nt from the 5’ end of *COX1*. The scale bar shows the distance in nucleotides, and the color code refers to the processing status of each read. E) Same as D) but for *ND1*. F) Quantification of all ribosome-protected fragments that align to the 5’ ends of mitochondrial genes. The plot shows the fraction of the unprocessed reads over all reads aligning to the 5’ end of the gene. Error bars show ranges of two replicates. G) Same as in F) but for 3’ ends. H) Fraction of 5’-end unprocessed mRNA from two replicate measurements of total RNA from FASTKD5 KD and matched control cells. I) Fraction of 5’-end unprocessed mRNA in total and ribosome-associated mRNA in the *FASTKD5* KD cells. Each dot or triangle represents data from one of two biological replicates. Triangles and darker-colored bars indicate the fraction of reads for which the read 5’-end starts in the tRNA closest to the 5’ end of the analyzed mRNA. J) Cartoon illustrating the effect of FASTKD5 depletion of mt-mRNA processing and ribosome association. All data in the figure are from HeLa cells except panels C, H, I, and J which are from HEK293T cells.

To test our hypothesis, we asked whether unprocessed mt-RNAs associate with the mitoribosome. We enriched for mitoribosomes and measured the full-length polyA+ translatome using direct RNA-seq. About 20% of *COX1* transcripts were unprocessed at the 5’-end in whole cell lysate, while less than 1% of transcripts were unprocessed in the mitoribosome (Figures 4C, and S4A, Table S3). Interestingly, an equally large proportion of *COX1* transcripts ended prematurely before the 3’-end, and this population was not strongly depleted in the mitoribosome, consistent with the observation that mRNAs lacking stop codons are nevertheless commonly translated by mitoribosomes^42,43^ (Figure 4C). For *ND1* and *ND2,* we found a smaller depletion of 5’-end unprocessed transcripts in the ribosome (Figure S4A). Upon closer inspection, we observed that the unprocessed transcripts were depleted of those with only a 5’ tRNA, and instead were unprocessed more upstream, and contained additional transcripts (Figure S4A). We wondered whether the *ND1* and *ND2* portions of the unprocessed transcripts were mitoribosome-associated as 3’ passengers without being actively translated. We analyzed mitoribosome profiling data to determine whether ribosome-protected fragments (RPF) contained unprocessed mt-mRNA (Figure 4D-G)^3^. We saw RPFs from several transcripts with unprocessed 3’-ends, but unprocessed 5’ ends were not found in RPFs aside from *COX3* as expected (Figures 4F-G, and S4B-D, and Table S6). Thus, translation initiation does not occur on 5’ unprocessed *ND1* and *ND2* mRNAs and their association with mitoribosomes is due to upstream RNA (Figure S4A).

The depletion of unprocessed RNA associated with mitoribosomes could be a consequence of fast RNA processing rather than a requirement for processed RNA in mitoribosome loading. We impaired 5’ processing to see if 5’-end unprocessed mRNAs would arrive to mitoribosomes by depleting cells of FASTKD5, a protein that participates in the 5’ processing of several mtDNA-encoded transcripts^50,51^. We found that FASTKD5 depletion leads to an increase in 5’-end unprocessed *ND1*, *ND2*, *COX1*, *COX3*, and *CYB* in total RNA, as previously reported (Figure 4F)^50,51^. Strikingly, 80% of *COX1* mRNA was unprocessed at the 5’-end in total RNA in the FASTKD5 KD cells, while less than 0.8% of the *COX1* reads in the ribosome were unprocessed at the 5’ end (Figures 4I, and S4E). Interestingly, for *COX1* a majority of mRNA 3’-ends now ended prematurely within the gene body in the FASTKD5 whole cell lysate. However, most of these transcripts were also unprocessed on the 5’ end and were not detected on the mitoribosome (Figure S4E, F). All 5’ unprocessed RNAs were underrepresented in mitoribosome purifications. As in wild-type cells, unprocessed mRNAs containing a 5’-end tRNA were strongly depleted in the mitoribosome, and the remaining unprocessed mRNAs had upstream rRNA or protein-coding genes (e.g. 5’ unprocessed *CYB* transcripts also retained the upstream *ND5* mRNA), explaining their mitoribosome association. In summary, mt-mRNA 5’-end processing was generally fast, and only *COX3* transcripts were detected in translating mitoribosomes with unprocessed 5’ ends (Figure 4F). Therefore, we conclude that 5’ processing is a translation prerequisite for all mt-mRNAs, except *COX3* (Figure 4J).

### Mitochondrial RNA turnover and translation efficiency are key predictors of protein synthesis levels

After determining rates for all major steps in the mitochondrial mRNA lifecycle, we sought to elucidate which processes represent the most important points of regulatory control in mitochondrial protein production. We developed a quantitative model that incorporated our measured RNA turnover rates (Figure 1D), transcription initiation and elongation rates (Figure 1K), and ribosome-association rates (Figure 4E, G) to predict protein synthesis levels calculated from ribosome profiling data^3^ (Figure 5A). The remaining variability we attributed to translation efficiency (TE = Ribosome profiling/RNA-abundance). RNA processing rates were not included as they are encompassed by ribosome-association rates. Applying this model, we found that RNA turnover had the largest impact, predicting 59.4% of the variability in protein synthesis levels, followed by translational efficiency, which explained 40.1% (Figure 5B). Transcription initiation and elongation did not contribute to the variation, and ribosome association slightly increased the predictive power by 0.5%. Thus, RNA turnover and translation efficiency are the main control points for mitochondrial gene expression.

**Figure 5.**
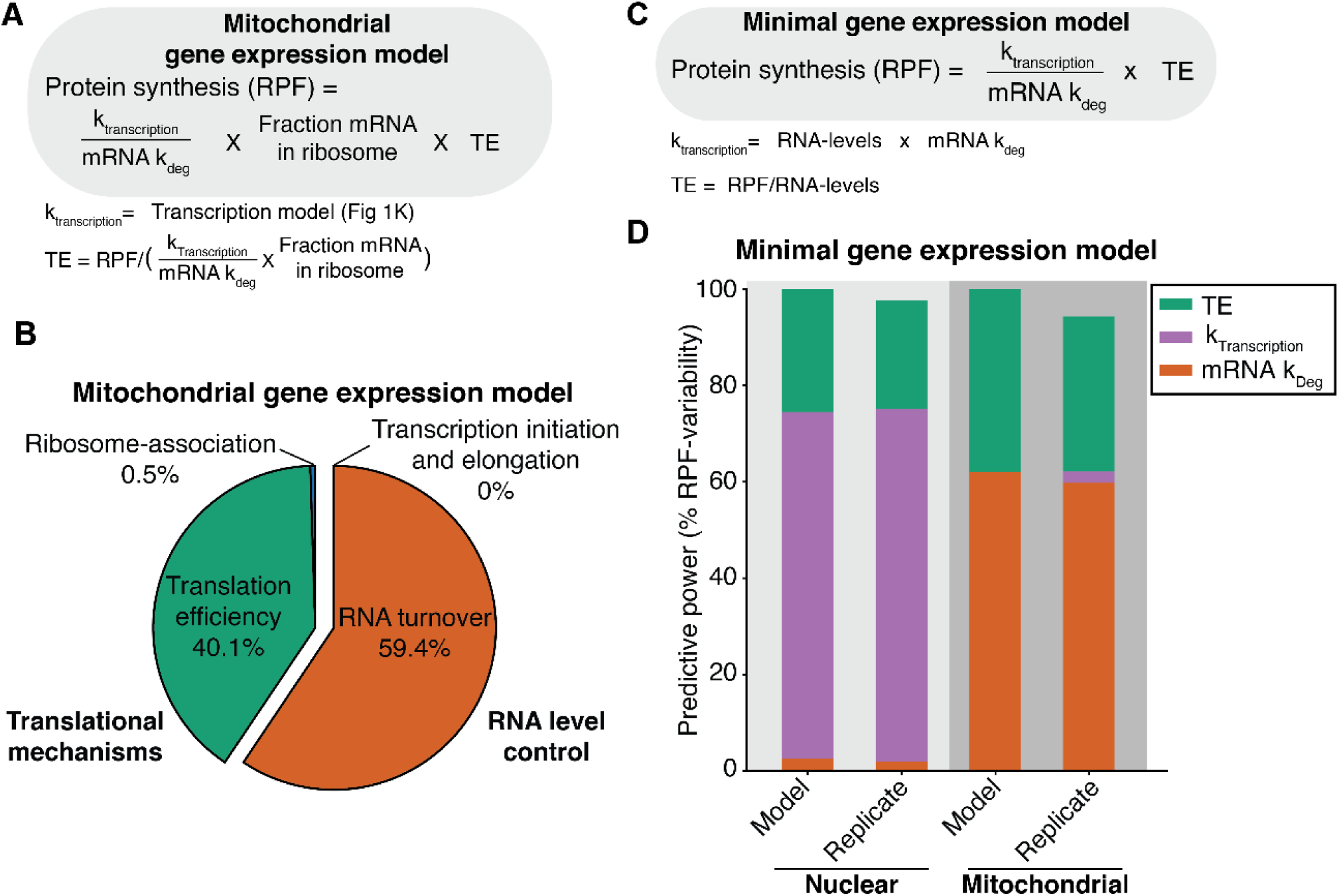
Mitochondrial gene expression is largely explained by RNA turnover and translation efficiencies. A) Quantitative gene expression model used in B) deploying the measured kinetic rates from this study. RPF: ribosome profiling, TE: translation efficiency, k_transcription_ = transcription rate, k_deg_ = turnover rate. B) Proportion of protein synthesis explained by different steps of gene expression in HeLa cells. RNA turnover and translation efficiency explain the vast majority of protein synthesis as determined by ribosome profiling, whereas ribosome association and transcription play minor roles. C) Minimal model of both nuclear and mitochondrial gene expression used in D). D) The ability of the nuclear and mitochondrial models to predict variability in HeLa ribosome profiling data for nuclear or mitochondrial-encoded OXPHOS genes, respectively. The “Model” by definition explains 100% of the variability, whereas the “Replicate" deploys the same estimated parameters to predict a replicate HeLa RPF dataset. Variability in nuclear gene expression is mostly explained by transcription levels whereas variability in mitochondrial gene expression is mostly explained by RNA degradation. TE was more predictive for mitochondrial genes than for nuclear-encoded genes.

### A minimal model predicts nuclear-and mitochondrial-encoded OXPHOS expression variation

We next sought to compare the impact of different gene expression steps within the nuclear and mitochondrial systems in setting OXPHOS synthesis levels (Figure 5C). We simplified our quantitative model to focus on the key points of gene regulation in each system. The model parameters were estimated independently for the mitochondrial and nuclear genes, using only mRNA turnover, transcription, and translation efficiencies (Figure 5C). Because the model was defined to explain 100% of the variability in the ribosome profiling data for a single biological replicate, we applied the model with the estimated parameters to predict RPF in a replicate dataset (Figure 5D); the coefficient of determination was >0.94. For nuclear genes, variability in the transcription rate, and not mRNA turnover, largely predicted variation in protein synthesis (Figure 5D). The reverse was true for mitochondrial genes: mRNA turnover, rather than transcription, was the predominant determinant (Figure 5D), consistent with our findings (Figure 1). Translation efficiencies were less predictive for nuclear-encoded OXPHOS genes than for mitochondrial-encoded genes, explaining ∼25% and ∼47% of the variation, respectively. In sum, these models combine the steps of the OXPHOS mRNA lifecycles (Figure 6A), illuminating the key stages of the contrasting kinetic regimes controlling nuclear and mitochondrial gene expression.

**Figure 6.**
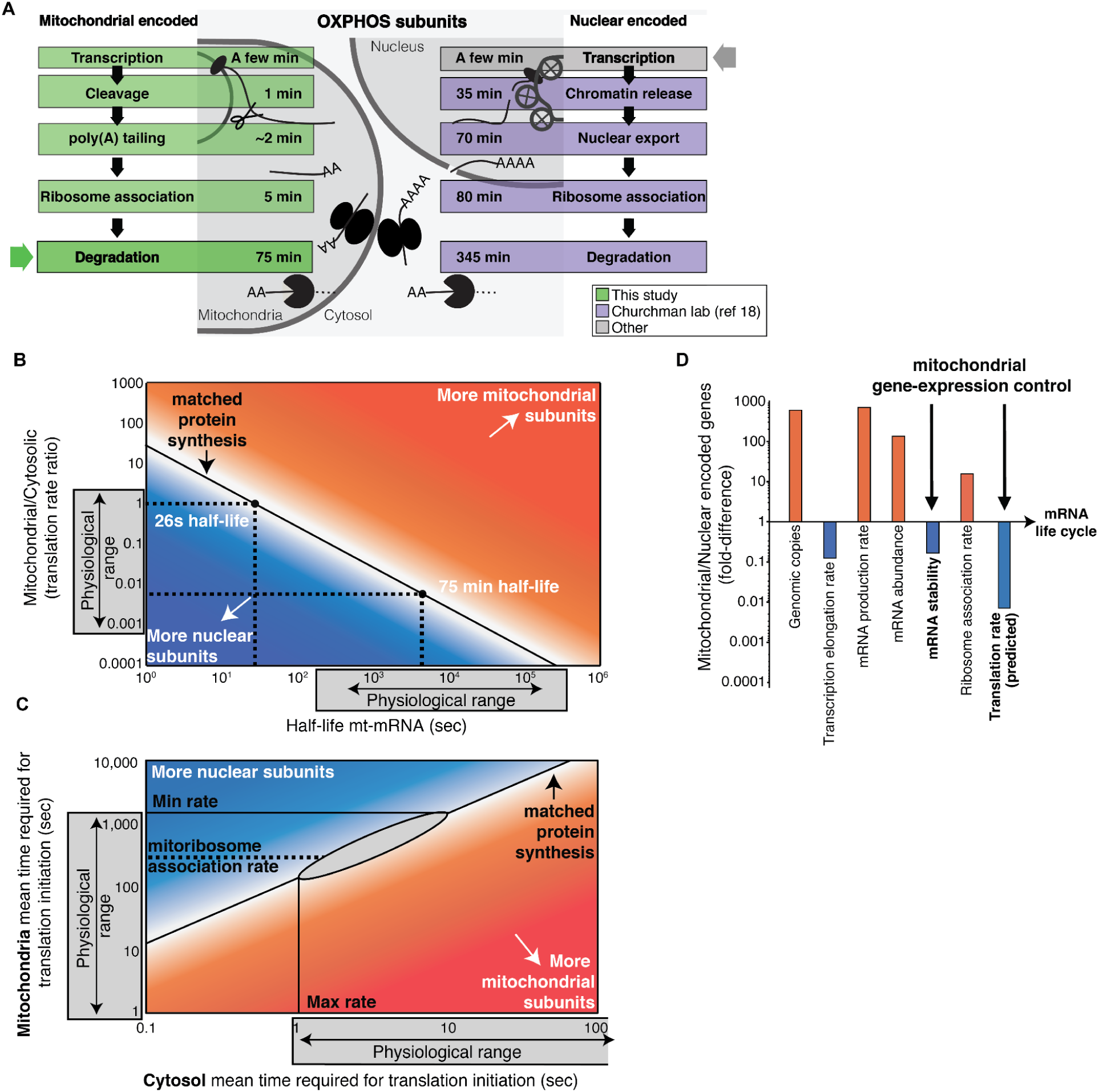
Comparison of the nuclear and mitochondrial gene expression layers. A) The life cycle of mitochondrial-encoded mRNA is on average much shorter than for nuclear-encoded OXPHOS mRNA. With the exception of transcription, the values are shown as time passed from production. Mitochondrial values are the median for HeLa cell in this study and nuclear-encoded mRNAs are the median from K562 cells from our earlier work^25^. The transcription parameters are based on estimates for how long it takes to produce one mRNA assuming a set transcription elongation rate. For mitochondrial mRNA, the estimate is based on our analysis and literature transcription elongation rates (range: 0.1-0.7 kb/min) and a median mRNA length of 1 kb. For nuclear OXPHOS encoding mRNA, the estimate assumes a median length of 7 kb and a transcription elongation rate of ∼2 kb/min^59^. We do not take into account promoter-proximal Pol II pausing. The colored arrows highlight the main gene regulatory level. B) Stoichiometric protein production of nuclear and mitochondrial-encoded OXPHOS subunits requires that the number mRNA multiplied by the translation rate constants (number of proteins per mRNA/h) is the same in both compartments. To match the cytosolic output, mitochondria can either adjust their translation rates or mRNA abundance. To reach the same mRNA abundance as the cytosol mt-mRNA half-lives need to be 26s on average (170-fold lower than 75 min which is the measured average). Likewise, a 170-fold lower translation rate in mitochondria compared to the cytosol would lead to matched synthesis. Our proposed physiological range (gray box) of mitochondrial mRNA half-lives are set to be higher than the ribosome association half-life, as to allow mRNA ever to be translated, and shorter than that of mt-rRNA. C) Translation initiation rates are usually considered rate-limiting for the overall translation rate^60^. Cytosolic translation initiation rate estimates vary widely^55,56^ and the maximum possible rate has been estimated to be one initiation event per second^56^. For matched translation outputs mitochondria translation initiation rates would be >100-fold slower. Here we set the slowest initiation rate to equal the shortest mRNA half-life. Our median mitoribosome association rate is indicated for reference. D) Order of magnitude analysis illustrates the great differences in gene expression between mitochondrial and nuclear-encoded genes. Values are from this study or from^25^ except for genomic copy numbers which are based on 1000 mtDNA copies compared to two nuclear chromosomes^61^. Translation rates are predicted as detailed in B).

## Discussion

### Highly polycistronic mitochondrial transcription drives the opposing kinetics of nuclear and mitochondrial gene expression

In this study, we developed a quantitative kinetic framework to analyze the human mitochondrial and nuclear-encoded transcriptomes, revealing many insights into OXPHOS gene regulation. Our measurements showed that RNA synthesis, processing, ribosome association, and degradation rates differed starkly between mitochondrial and nuclear-encoded OXPHOS genes (Figure 6A). Why do these differences arise, and how are they overcome? Nascent mitochondrial RNA is polycistronic, producing mRNAs at similar levels to non-coding RNAs, which must accumulate highly to support mitochondrial translation. As mt-rRNAs are positioned immediately after the heavy-strand transcription start site (Figure 1A), a separate mt-rRNA transcription module has been proposed to explain the high abundance of rRNA compared to mitochondrial mRNA^52^. Here instead, we found that the production rates of mt-mRNA were the same as those of mt-rRNA while steady-state RNA levels varied widely (Figure 1E, F). Our heavy-strand polycistronic transcription model explained these observations by highlighting the importance of gene-specific degradation rates (Figure 1K). In contrast, nuclear-encoded mRNA levels are predominantly regulated by gene-specific transcription start sites. The high co-production of mt-rRNA and mt-mRNA also necessitates the 5-fold more rapid mt-mRNA turnover rates than nuclear-encoded OXPHOS mRNA. Because if mt-mRNA were as stable as mt-rRNA or nuclear-encoded OXPHOS mRNA, mt-mRNA would represent 70% or 25% of the cellular mRNA pool, respectively, rather than the 6% it consists of actually. Thus, the different degradation kinetics of the mitochondrial and nuclear gene expression systems are necessary to support the highly polycistronic nature of human mitochondrial pre-RNA.

### Slow mitochondrial translation is required to balance OXPHOS gene expression

RNA degradation is insufficient to balance the nuclear and mitochondrial transcriptomes; we observed a ∼170-fold difference in OXPHOS transcript abundance between the mitochondrial-and nuclear-encoded OXPHOS mRNAs (Figure 1E). To achieve a balanced OXPHOS transcriptome by RNA degradation alone, RNA half-lives would have to be around 30 seconds (Figure 6B), which would be too short for even one round of translation. Thus, a scaling problem emerges if we assume the stoichiometric accumulation of OXPHOS complex subunits. Because we found mitochondrial transcripts to be predominantly associated with ribosomes (Figure 3), the mitochondrial translation rate would need to be >100-fold slower than cytosolic translation (Figure 6B). Alternatively, mitochondrial-encoded proteins could be turned over much faster than their nuclear-encoded counterparts; however, mitochondrial-encoded proteins are stable with half-lives much longer than the cell doubling time^53^. Thus, a low mitochondrial translation rate is the most likely explanation and is consistent with higher mitoribosome levels per OXPHOS protein compared to cytoribosomes^54^. Estimates of cytosolic translation rates in HeLa cells range from 180-280 proteins/mRNA/h by single mRNA translation kinetics measurements^55^ and up to 10-fold higher estimates by modeling on proteomic, transcriptomic, and translatomic data sets^56,57^. Accordingly, cytosolic translation initiation times then range between 1-20 seconds^55,56^. The equivalent numbers for mitochondria would need to be 1-25 proteins/mRNA/h and translation initiation times of 140-3600 seconds, in the range of our ribosome association rates which average 380 seconds (Figure 6C). Since some mt-mRNAs exist for less than an hour, many mt-mRNAs would be translated only a few times. Alternatively, mitoribosome stalling while awaiting the arrival of nuclear-encoded subunits has been proposed to enable the association of mitochondrial-encoded COX1 and nuclear-encoded COX4 during Complex IV assembly, which would also result in slower mitoribosome translation^58^. In sum, we propose that slower mitoribosome translation promotes mito-nuclear balance by scaling mitochondrial protein production to match cytosolic OXPHOS synthesis, which may simultaneously serve in the coordination of OXPHOS assembly.

OXPHOS biogenesis is a particularly complicated cellular production process as it requires the coordination of two gene expression systems housed in separate compartments and, as we show here, are opposed at nearly every step. Moreover, it is precarious; misassembled respiratory chains lead to toxic consequences^62–64^. Thus, the balanced assembly of OXPHOS complexes is a triumph of the eukaryotic cell, providing an energetic evolutionary advantage that fostered genome complexity and, ultimately, multicellularity^65,66^. However, considering the role of mitochondrial malfunction in numerous disease and aging phenotypes^5,67–70^, maintaining mitonuclear balance may represent an Achilles heel of the animal cell. Our quantitative and kinetic framework for studying mitochondrial and nuclear gene expression provides a critical foundation for understanding how the cell navigates the mito-nuclear challenge in healthy cells and how disease comes about when it fails.

## Acknowledgments

We thank Anna Wredenberg (Karolinska Institute), Christoph Freyer (Karolinska Institute), Jake Bridgers, and Chantal Guegler for their careful review of the manuscript. We thank members of the Churchman lab, Matthias Selbach (Max Delbrück Center for Molecular Medicine), and Karl McShane (Malmö Stad) for fruitful discussions. We thank Ari Sarfatis for helping develop the data analysis pipeline; the Biopolymers Core Facility at Harvard Medical School for sequencing services and the Boston Children’s Hospital Molecular Genetics Core for NanoStrings services.

## Availability of data and materials

Raw and processed data will be available from GEO. Code for analysis of all data will be made publicly available at http://github.com/churchmanlab/mitoRNA_kinetics.

## Methods

### Cell culture

HeLa cells were grown in DMEM containing glucose and pyruvate (ThermoFisher, 11995073), supplemented with 10% FBS (Thermo Fisher, CAT A3160402.). LRPPRC KO and matched WT HEK293T cells from^3^ were cultured in DMEM supplemented with 20 μg/mL of uridine (Sigma, CAT U3750), 3 mM sodium formate (Sigma CAT 247596), 10% FBS, 1 mM GlutaMAX (Thermo Fisher Scientific, CAT 35050061) and 1 mM sodium pyruvate (Thermo Fisher Scientific, CAT 11360070). Human myoblasts from anonymous healthy donors were kindly provided by Dr. Brendan Battersby (Institute of Biotechnology, University of Helsinki). Myoblasts were grown in Human Skeletal Muscle Cell Media with the provided growth supplement (HSkMC Growth Medium Kit, Cell Applications, 151K-500). For differentiation into myotubes, the media was replaced with DMEM (ThermoFisher, 11995073) with 2% heat-inactivated horse serum and 0.4 ug/mL dexamethasone. The differentiation media was replaced every two days.

### Creation of knock-down cells line

CRISPR mediated knock-outs were performed by transducing HEK293 cells with LentiCRISPRV2 (Addgene #52961) lentivirus expressing sgRNAs targeting genes of interest (FASTKD5_sg1: CACCGCATATGGTGCTTGACGTACT; FASTKD5_sg2: CACCGGCTGTGAACACGGTATTC). Transductions were performed by addition of lentivirus containing supernatants to 80% confluent 293T cell cultures in the presence of 8 ug/mL polybrene for 16-24 hr. Upon which the media was replaced with fresh DMEM for 24h, followed by selection with puromycin (2 ug/mL) for 72h.

### 4sU labeling

HeLa or HEK293T Cells were grown to 70-80% confluency and then pulsed by either 50, 100, or 500uM final concentration of 4-Thiouridine (4sU, Sigma T4509) in pre-conditioned cell culture media under standard cell culture conditions. For mitoribosome-IP experiments, one 15 cm plate of cells was used per time point while for all other experiments, one 10 cm plate was used per time point.

### Cell lysis and RNA extraction

Labeled cells were washed in ice-cold PBS (ThermoFisher) before being directly scraped in Trizol LS (ThermoFisher, 10296010). RNA extraction was performed using Trizol LS according to the manufacturer’s protocol, except with the addition of DTT (Sigma-Aldrich) at a final concentration of 0.2 mM DTT in the isopropanol.

Undifferentiated myoblasts (day 0) and differentiated myotubes (day 7) were collected and lysed in 1 mL of Qiazol (Qiagen). RNA from undifferentiated and differentiated myoblasts was extracted using the Qiagen miRNeasy kit (Qiagen) according to the manufacturer’s protocol.

### mtPolysome fractionation, mitoribosome-IP, and RNA extraction from mitoribosome-IP samples

After labeling cells were quickly washed in ice-cold PBS before being scraped in mt-polysome lysis buffer (0.25% Lauryl Maltoside, 10mM Tris pH 7.5, 0.5mM DTT, 20 mM magnesium chloride, 50 mM ammonium chloride 1× EDTA-free protease inhibitor cocktail (Roche)). Cell lysates were dounced 7 times in a 1mL dounce with the tight piston and then flash frozen. 1mL of thawed lysate was clarified by spinning twice at 10,000 rcf. To avoid contamination of nascent RNA still attached to mtDNA into the higher molecular weight sucrose gradient fractions we digested all DNA by 150 units of DNAse I (NEB) in the presence of superaseIn (ThermoFisher) and 0.5mM calcium chloride at room temperature for 1 h. To isolate mitoribosomes, lysates were loaded on 10–50% linear sucrose gradients and centrifuged in a Beckman ultra-centrifuge at 40,000 RPM for 3 h at 4°C using a SW41Ti rotor. Some of the input for the sucrose gradient fractionation was kept and treated as a standard sample described above. Gradients were mixed and fractionated using a BioComp instrument. To identify the mitoribosome containing fractions we western blotted for Mrpl12 (Proteintech 14795-1-AP) and Mrps18B (Proteintech 16139-1-AP) as described below. Monosomes and polysomes containing fractions were pooled and the mitoribosomes were immunoprecipitated out of the pooled fractions. For the immunoprecipitation, MRPL12 antibodies were conjugated to Protein A dynabeads (ThermoFisher) for 1 h in mt-polysome lysis buffer. After washing the beads, lysates were added and incubated for 3 h at 4C. After 3 h the supernatant was removed and the beads were washed three times in mt-polysome lysis buffer before the mitoribosomes were eluted in 0.2% SDS, 100 mM NaCl, 10 mM Tris pH 7.5, 1X EDTA-free protease inhibitor cocktail and SuperaseIn. IP efficiency was confirmed by western blotting as described below. RNA was extracted from the eluates using Trizol LS as described above and then further cleaned by using 1.8x volume of RNAClean XP beads (Beckman Coulter A63987) and washed in 80% ethanol.

When collecting RNA, from NT or FASTKD5 KD cells, destined for direct RNA-sequencing on the nanopore two gradients per condition were used as input for the IP and the final clean-up step using RNAClean XP beads was omitted.

### Western blotting

Samples were mixed with 4 x LDS sample buffer (ThermoFisher NP0007) and 0.1M DTT. Samples were loaded onto a 4-12% Bis-Tris gel (Invitrogen NP0321BOX) in 1x MOPs buffer and run at 160V for 1 hour. The gel was transferred to a nitrocellulose membrane using the wet transfer method in 1x transfer buffer (25mM Tris base, 192mM glycine, 20% methanol) at 400mA for 75 minutes at 4°C. The membrane was blocked in a blocking buffer (5% non-fat milk powder in 1x Tris buffer saline with 0.1% Tween 20) for at least 60 minutes. Primary antibodies were diluted in blocking buffer and incubated with membranes overnight at 4°C. Membranes were washed 3 x 20 minutes with 1 x TBST, incubated for 1 hour at 25°C with secondary HRP-conjugated antibodies against rabbit IgG (Cell Signaling Technology, 7074S), washed again 3x 20 minutes with 1x TBST, and developed using ECL Western Blotting Detection Reagents (Cytiva) and Amersham Hyperfilm ECL (Cytiva) using an automated M35 X-OMAT Processor developer (Kodak).

### TimeLapse-sequencing

Extracted 4sU-labeled RNA from HeLa or HEK 293T cells were treated by TimeLapse-chemistry as described in^21,25^. In short, 2ug of RNA was treated with 0.1M sodium acetate pH 5.2, 4mM EDTA, 5.2% 2,2,2-trifluoroethylamine, and 10mM sodium periodate at 45°C for 1 hour. RNA was then cleaned using equal volume or RNAClean XP beads (Beckman Coulter A63987) and washed in 80% ethanol. The cleaned RNA was reduced in 0.1M DTT, in 0.1M Tris pH 7.5, 1M NaCl, and 0.01M EDTA for 30 min at 37 C before once again being cleaned by RNAClean XP beads. Sequencing libraries from treated RNA were created using the SMARTer Stranded Total RNA HI Mammalian kit (Takara 634873) following the manufacturer’s instructions. Libraries were sequenced on the NextSeq (Illumina, San Diego, CA) by the Biopolymers Facility at Harvard Medical School.

For the mitoribosome-IP the rRNA depletion step was left out for the IP:ed RNA as their cytosolic rRNA was already depleted by the IP step. In addition, the SMARTer RNA unique dual index kits (Takara 634452) were used instead of the primers that came with the SMARTer Stranded Total RNA HI Mammalian kit to allow for sequencing on the NovaSeq (Illumina, San Diego, CA) by the Biopolymers Facility at Harvard Medical School.

### Creation of SNP-masked cell-line-specific reference genomes

To create cell-line-specific genomic single nucleotide polymorphism-corrected reference sequences, we used reads from samples without 4sU treatment. Reads were first aligned to the reference hg38 genome with STAR using parameters --outFilterMultimapNmax 100 --outFilterMismatchNoverLmax 0.09 --outFilterMismatchNmax 15 --outFilterMatchNminOverLread 0.66 --outFilterScoreMinOverLread 0.66 --outFilterMultimapScoreRange 0 --outFilterMismatchNoverReadLmax 1. Variants were then called with BCFtools (Li 2011) ‘mpileup’ and ‘call’ using two bam files as input. The resulting variant call file (VCF) was then split into a file with INDEL records only and a file without INDEL records (substitutions only). The "no INDEL’’ VCF was further split by frequency of substitution: loci covered by >= 5 reads and with a variant frequency >75% to a single alternate base were assigned the alternate base; loci with variants with an ambiguous alternate base were masked by "N" assignment. The reference FASTA was modified for these non-INDEL substitutions using GATK (McKenna et al 2010) FastaAlternateReferenceMaker. Finally, rf2m (https://github.com/LaboratorioBioinformatica/rf2m) was used with the INDEL-only VCF file to further modify the FASTA genome reference as well as the corresponding GTF annotation file.

### Read Alignments

First, fastq files shared by the sequencing facility were concatenated to combine data across the sequencer lanes. The adapter sequences were trimmed from both of the paired-end reads using Cutadapt v2.5 (Martin. EMBnet, 2011). Three more nucleotides were trimmed from the 5 prime end of the fragment to remove template switch bases, and five more from the 3 prime end to remove mismatches associated with random hexamer priming that result from the SMARTer Stranded library preparation. Read alignment was done using STAR 2.7.3 to cell-line-specific genomic reference sequences (see above) with different parameters for experiments with high and low 4sU treatment. For 500 uM 4sU experiments, parameters were --outFilterMismatchNmax 90 --outFilterMismatchNoverLmax 0.3 –outFilterMatchNminOverLread 0.2 --outFilterScoreMinOverLread 0.2. For 50 uM 4sU experiment, parameters were --outFilterMismatchNmax 10 --outFilterMismatchNoverLmax 0.05 --outFilterMatchNminOverLread 0.66 --outFilterScoreMinOverLread 0.66 --outFilterMultimapScoreRange 0.

### T to C mismatch counting

For each sample, a subset of mitochondrial-aligned reads was used to determine the rate of T to C mismatches. Unless otherwise specified, this subset included all mitochondrial reads except for those aligning to MT-rRNA. For this analysis, reads were first filtered to remove non-primary alignments, reads with unmapped mates, and reads that map to more than 4 positions, and soft-clipped bases were removed. We then determined the number of T to C mismatches and the total number of T nts across each fragment (considering both read1 and read2 in each pair) using custom scripts from^25^. Two-dimensional distributions were generated containing the number of fragments with *n* Ts and *k* T to C mismatches.

### Binomial mixture model to calculate T to C conversion rates

To estimate the 4sU-induced as well as background T to C conversion rates, we used a binomial mixture model we previously developed ^25^ to handle samples with low 4sU concentrations. In short, the binomial mixture model first estimates two background T to C conversion rates (*p*_*E*1_ and *p*_*E*2_) and a global fraction (π_*E*_) parameter, with *Binom* a binomial distribution, given the measured *n* Ts and *k* T to C mismatches in the unlabeled control sample. Giving the background T to C conversion rates as follows: *P*_*BG*_(*k*, *n*) = π_*E*_ *Binom*(*k*, *n*, *p*_*E*1_) + (1 − π_*E*_)*Binom*(*k*, *n*, *p*_*E*2_). The three parameters in this model were fitted to the above T to C distributions using linear regression (Python v3.7.4, package lmfit v1.0.2, function minimize). We tested if our binomial mixture error model better fitted the T to C conversions of the untreated samples using the Akaike Information Criterion and adjusted our background model to include one or two background rates depending on the test outcome^71^. The 4sU sample T to C distributions were then modeled as the 4sU-induced T to C conversions (*p*_*c*_) plus the background population *P*_*BG*_ (*k*, *n*) once again with a global fraction (π_*c*_) parameter: *P*(*k*, *n*) = π_*c*_ *Binom*(*k*, *n*, *p*_*c*_) + (1 − π_*c*_)*P*_*BG*_ (*k*, *n*). We note that it is necessary to calculate the T to C conversion rate from nuclear and mitochondrial encoded transcripts separately as the mitochondrial 4sU-incorporation rate is significantly lower than for nuclear-encoded genes. In experiments deploying high (500uM) 4sU concentration with no Uridine supplemented to the media, we had robust T to C conversion rates >1.5% and directly used GrandSlam to estimate conversion rates. The T to C conversion rates in the mitoribosome IP and input samples as well as samples from HEK293T cells were calculated from all reads aligning to the mitochondrial genome except reads aligning to COX1 and RNR2.

### T to C conversion rate estimates for cases with very low 4sU incorporation

In three cases we ended up with T to C conversion rates lower than 0.6%: the earliest (15 min) time point for the HeLa mitoribosome IP experiments deploying 100uM 4sU (Figure 3D, E), all the time points in the HEK293T mitoribosome IP experiments (Figure S3F) and for all the time points in the LRRPRC KO timecourse. In these cases, our binomial method failed and we instead calculated the T to C conversion rate based on the assumption that the ribosomal RNA would be turned over consistently. Practically, we set the T to C conversion rate such that it resulted in the fraction of new RNR2 being the same as at the equivalent time point in the high confidence time course experiments in Figure 1 or the WT cells in the case of the LRPPRC KO. Importantly, the RNR2 half-lives were estimated to be the same in the low T to C conversion experiments as in the high T to C conversion experiments, but with a lower confidence. In the IP we set the T to C conversion rate so that the fraction of new RNR2 was lower than 0.5%, to allow for some noise. This method gave similar results to the mitoribosome IP experiment with higher 4sU levels where we did not use any corrections (Figure 3E).

### GRAND-SLAM parameters

Alignment files (.bam) containing all mitochondrial-aligned reads were converted into a .cit file using GRAND-SLAM v2.0.5f^24^. The no-4sU (0 min) alignment file was copied and both copies (“no4sU” and “0m”) were included in the GRAND-SLAM analysis so that the “no4sU” sample would be used for the background T to C mismatch rate (p_E_) and the sample details for the “0m” sample would also be output. GRAND-SLAM was run twice: first for .cit file creation and background rate determination, and a second time after modifying the T to C mismatch rates (p_C_) according to the output of the binomial mixture model. Before the second run, the *.tsv and *ext.tsv files were removed from the output directory and the *rates.tsv file modified to add in new rates for single_new and double_new. For both runs the parameters used included -full -mode All -strandness Sense -snpConv 0.4 -trim5p 10 -trim3p 5.

### Cell doubling correction of fraction new values

The fraction new RNA values for HeLa cells was corrected for dilution by cell doubling. The cell doubling time was estimated by growing HeLa cells under standard cell culture conditions to 70% confluency (0h). At 0h, 12h, and 24h, five replicates of 10cm plates were washed in PBS, and cells were collected by trypsinization. Cells were mixed with trypan blue and counted using an automated cell counter (Countess, Invitrogen). Cell doubling time was estimated to be 26.5 h by linear regression of the log2-fold change in cell number over time. The fraction new values were corrected as follows:

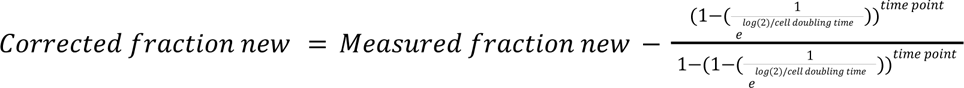

### Kinetic modeling of mtRNA degradation

Based on the maximum a posteriori (MAP) values obtained from GRAND-SLAM for each time point of the 4sU pulse, we tested two simple models to estimate the degradation rates from the time t dependence of newly synthesized <u>T</u>otal mtRNA *T*(*t*) ^24,25,72^.

Model 1: exponential (one-state) decay - RNA is produced with a transcription (<u>p</u>roduction) rate *k*_*p*_ and eventually degraded at the rate *k*_*deg*_ (Figure S1F): 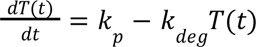. To solve this ordinary differential equation analytically as described previously ^24,25,72^, we set all newly synthesized RNA levels to zero at t=0, i.e. before any 4sU pulse as the boundary condition. Next, the integrating factor method was used to obtain the solution: *T*(*k*_*p*_, *k*_*deg*_, *t*). Since the observed quantities are fraction of new RNA, rather than RNA levels we derived the model fraction of newly synthesized RNA: 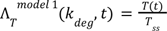, with 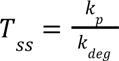 the <u>s</u>teady <u>s</u>tate levels of all (newly synthesized and pre-existing unlabeled) RNA, which equals the steady state solution to the ODE: 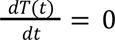. Both *T*_*ss*_ and *T*(*t*) are linear in *k*_*p*_, so Λ_*T*_ ^*model* 1^ (*k*_*deg*_, *t*) no longer depends on *k*_*p*_. So, we arrive at our solution:

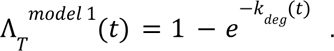

Model 2: non-exponential (two-state) decay - RNA first assumes state 1 *S*_1_ (*t*) upon which it can either undergo exponential decay at the rate *k*_*d*1_ or it can transition to state 2 *S*_2_ (*t*) with rate *k*_*r*_ from which it exponentially degrades with rate *k*_*d*2_ (Figure S1F):

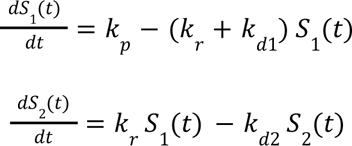

Here, the total amount of mtRNA is then given by the sum of both states: *T*(*t*) = *S*_1_ (*t*) + *S*_2_ (*t*). Using the same procedure as described above for the 1 state model, the fraction new total RNA then becomes (equation 1)^72^:

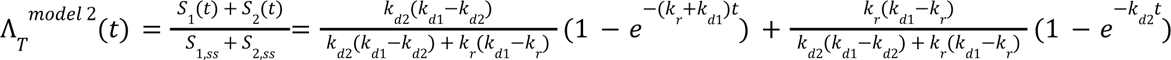

To estimate the degradation parameters in model 1 (one parameter) and model 2 (three parameters), we used a non-linear least square fitting framework (Python v3.7.4, package lmfit v1.0.2, function minimize). Note, for all experiments we directly used the MAP values from GRAND-SLAM for the modeling.

To select the model to make predictions about the RNA decay parameters, we used the Akaike information criterion 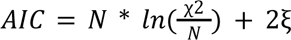, where N denotes the number of data points, χ^2^ is residual square sum and ξ is the number of variable parameters. To further estimate which model has a higher probability we can define model weights to define the probability of model 1 over model 2. The difference in AIC for a given model *i* is given by 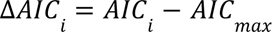, and AIC_min_ is the minimum AIC across all models. Based on this Akaike weights are estimated by 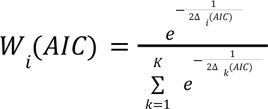, where K is the number of models. Since we are comparing two models we defined the normalized probability as π by 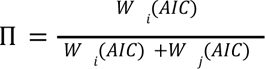

### Kinetic modeling of mtRNA association to the mitoribosome

To estimate the ribosome association rate, we reused the two-state model as described above, with the free mtRNA state, i.e. not bound to the ribosome, *F*(*t*) = *S*_1_ (*t*) and the ribosome-bound mtRNA state *RB*(*t*) = *S*_2_ (*t*). The ribosome association rate is then the transfer rate *k*_*r*_ from the free state to the ribosome-bound state. In addition to the total RNA (equation 1 above), we now also fit the experimentally observed fraction of newly synthesized ribosome bound mtRNA:

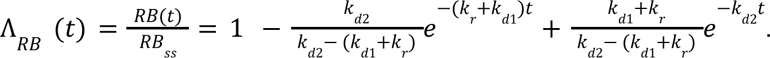

We can further simplify these equations by making the assumption that *k*_*deg*_ = *k*_*d*1_ = *k*_*d*2_:

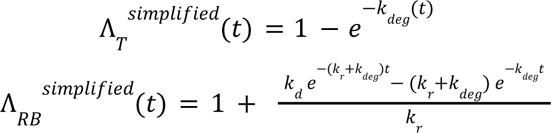

To estimate the degradation and mitoribosome association rates in the simplified model (two parameters) and full model (three parameters), we used a non-linear least square fitting framework (Python v3.7.4, package lmfit v1.0.2, function minimize) to fit the total and ribosome-bound mtRNA. To select the model to make predictions about the mtRNA kinetic parameters, we used the Akaike information criterion as discussed above.

Lastly, to calculate the fraction of nuclear-encoded mRNA that was bound by the cytosolic polysome, we derived the following equation, with nuclear residence *k*_*nuc*_, cytoplasmic turnover *k*_*cyto*_ and cyto-polysome entry rates *k*_*r*_ as determined in^25^:

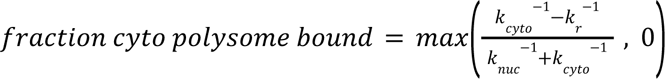

### Transcription elongation model

In order to model the mitochondrial transcription elongation process, we first processed the Nanopore sequencing reads according to the assumption that total RNA consists of nascent and mature RNAs. Reads with a 3 prime end within a 100nt window of annotated mature RNA 3 prime ends contribute to the mature RNA fraction with coverage across the whole of the gene. Reads with ‘3 ends within the annotated genes, but outside of the 3 prime end window contribute to the nascent RNA fraction with coverage across the whole genome up to the from the single TSS in the mitochondrial genome, i.e. the HSP2 3 prime end site. The sum of nascent and mature RNA fractions makes up the total RNA coverage across the genome. The model mature RNA fractions equal 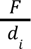 and the model nascent RNA fraction equals 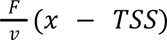, with x the distance (unit: nt) downstream of the TSS, as previously described in^28,29^. *F* is the transcriptional initiation (<u>F</u>iring) rate, *ν* the MT Polymerase elongation rate, and d_i_ the mature RNA degradation rate, specific to gene i, as estimated above through TimeLapse sequencing. The initiation rate F can then be estimated from the observed mature RNA counts from each gene i: mature RNA_i_ = F_i_ / d_i_. This leads to an ensemble of F estimates: {*F*_*i*_ }. Since RNR2 is the most upstream and highly expressed, we used this “best estimate”: 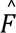. We also calculate the *F*_*min*_ = *min*_*i* ∈*genes*_ {*F*_*i*_} and *F*_*max*_ = *max*_*i* ∈*genes*_{*F*_*i*_} as the estimate bounds. To estimate *ν*, we use linear regression to fit the nascent RNA fraction model, whilst utilizing our {*F*_*i*_}, to get an elongation rate ensemble {*ν*_*i*_} (Python package: lmfit, function: minimize, velo_seed = 3.81 * 60, equalling the previous estimate by^33^, velo_max = 100*1000 #units: nt / min) from RNR2 3 prime end to the 3 prime end of the genome. 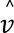 is then the elongation rate paired with 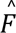, and *ν*_*max*_ = *max*_*i* ∈*genes*_ {*ν*_i_}, likewise for *ν*_*min*_. Altogether, our estimates are then used to predict total RNA coverage with (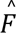, 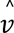) as best estimate (red lines) and (*F*_*min*_, *ν*_*min*_) and (*F*_*max*_, *ν*_*max*_) as error bands (Figure 1H).

### Ribosome footprinting analysis

Mitochondrial ribosome profiling was initially processed as previously described^3^. To analyze the 5 prime and 3 prime-end processing status of ribosome-loaded mt-mRNA, the position of both the 5 prime and 3 prime ends of each read were recorded. Soft-clipped bases were ignored for recording positions but flagged if they were determined to come from a 3 prime poly(A) tail (>80% of soft-clipped bases are A). A read was considered processed if its terminus is within two nucleotides of the processed transcript terminus. A read was considered unprocessed if it spans at least three nucleotides beyond a processed transcript terminus and was not flagged as containing a poly(A) tail.

### MitoStrings experiments

mt-RNA turnover in HeLa cells was determined by measuring the decrease of the old/unlabeled mtRNA after a 4sU-pulse as described in^25^. In short, HeLa cells were labeled with 500uM 4sU for 0, 60, or 120 min and cells were lysed in Trizol LS and RNA was extracted according to the manufacturer’s protocol. A spike-in of *in vitro* transcribed ERCC-000148 was added to all the cell lysates during the RNA-extraction step. The RNA was denatured at 60C for ten minutes before being biotinylated by the addition of 5ug/ml biotin-MTS (Biotium) in 20% dimethylformamide (Sigma), 20mM Hepes pH 7.4 and 1 mM EDTA for 30 min at room temperature. Free biotin was removed using phase-lock heavy gel tubes (5prime) following the manufacturer’s instructions. Biotinylated RNA was removed by incubation with streptavidin beads from the uMACS Streptavidin kit (Miltenyi Biotec) for 15 min at room temp. Beads and RNA were loaded onto a uMacs column. Beads were washed with 100mM Tris pH 7.5, 10mM EDTA, 1M NaCl, 0.1% Tween 20, and unlabeled RNA was collected in the flowthrough. RNA in the flow-through was purified using the miRNeasy Nano kit following the manufacturer’s instructions including the DNAse treatment step (Qiagen). 30 ng of RNA was incubated for 16h at 67°C with the XT Tagset-24 (NanoString Technologies) and with DNA-probes specific for mitochondrial RNA (Original Mitostring-probes^26^ were modified as in^73^) in hybridization buffer (NanoString Technologies) according to the manufacturer’s protocol before being loaded onto a nCounter Sprint Cartridge and quantified using the nCounter SPRINT Profiler (NanoString Technologies) at the Boston Children’s Hospital Molecular Genetics Core.

For direct measurements of processing kinetics HeLa cells were labeled with 500uM 4sU for 0, 1.875, 3.75, 7.5, 15, 30, 60, 120, or 240 min. RNA was extracted as described above with the addition of a second *in vitro* transcribed 4sU-labeled (10% of UTP was exchanged for 4sUTP in the T7 RNA polymerase *in vitro* reaction (NEB)) spike-in ERCC-00136. Biotinylation and streptavidin-bead binding were performed as above with the exception that in this case the flowthrough was discarded and bead-bound nascent RNA was eluted by incubating the beads at 2 x 5 min at 65C in 100mM TCEP (Thermo Scientific), 10mM EDTA, 500mM Tris at a final pH7. Eluted RNA was purified, hybridized with Mitostring probes, and analyzed on a nCounter Sprint Profiler as above.

### Direct nanopore RNA sequencing

RNA was extracted as described above using Trizol LS and chloroform. Extracted RNA was rRNA-depleted by hybridization with a custom set of biotinylated-DNA probes complementary only to cytosolic rRNA (riboPOOL, siTOOLS). In short, 6ug of RNA was mixed with 5uM riboPOOL probes in 20 uL of 2.5mM Tris-HCl (pH 7.5), 0.25mM EDTA, and 500uM NaCl and hybridized by incubating at 68°C for 10 min followed by a slow cooling step decreasing the temperature to 37°C. 20uL of hybridized samples were mixed with Dynabeads MyOne Streptavidin C1 beads (ThermoFisher) in 80uL of 5mM Tris-HCl (pH 7.5), 0.5mM EDTA and 1M NaCl. Dynabeads had been pretreated with sodium hydroxide according to the manufacturer’s instructions to minimize RNase contamination. The sample and bead mix was incubated at 37°C for 15 min followed by a 5 min step at 50°C before beads were collected on a magnet and the supernatant was transferred to a new tube with beads and the incubation steps were repeated. RNA collected in the supernatant from the second round of bead depletion was cleaned up by isopropanol precipitation.

To include both non-polyadenylated and polyadenylated transcripts, 500 ng of rRNA-depleted RNA was polyadenylated *in vitro* using *E. coli* poly(A) polymerase as outlined in^74^. As an alternative strategy, we ligated a DNA linker (/5rApp/(N)6CTGTAGGCACCATCAAT, IDT) to the 3 prime-end of total RNA as previously described^75^. Briefly, 1 ug of total RNA was denatured for 2 minutes at 80°C and incubated with 500 ng DNA linker, 8 uL 50% PEG8000, 2 uL DMSO, 2 uL of T4 RNA Ligase Buffer (NEB) and 1 uL T4 RNA Ligase 2 (made in-house) in a 20 uL reaction for 16 hours at 16°C. The RNA was purified with the Zymo RNA Clean & Concentrator Kit, including the optional removal of molecules < 200 nt to exclude the leftover linker. Direct RNA library preparation was performed using the kit SQK-RNA002 (Oxford Nanopore Technologies) with 500 ng of poly(A)-tailed or 3 prime-end ligated RNA according to manufacturer’s instructions with the following exceptions: the RCS was omitted and replaced with 0.5 uL water and the ligation of the reverse transcription adapter (RTA) was performed for 15 minutes. For 3 prime-end ligated samples, the standard RTA was replaced by a custom adapter that was generated by annealing two oligonucleotides (IDT): /5PHOS/GGCTTCTTCTTGCTCTTAGGTAGTAGGTTC (ONT_oligoA) and GAGGCGAGCGGTCAATTTTCCTAAGAGCAAGAAGAAGCC<u>GATTGATGGT</u> (ONT_oligoB_linker), where the underlined bases are complementary to the DNA linker. Oligonucleotides were diluted to 1.4 μM in 10 mM Tris-HCl pH 7.5, 50 mM NaCl and annealed by heating to 95°C and slowly cooling to room temperature.

Poly(A)+ RNA from myoblasts, HeLa, and HEK293 (NT and FASTKD5 sgRNAs) samples were purified using the Dynabeads mRNA purification kit (ThermoFisher, 61006) according to the manufacturer’s instructions, starting with up to 75 ug of total RNA. Direct RNA library preparation was performed as described above with 500-700 ng of poly(A)+ RNA. Direct total poly(A)+ RNA sequencing data from K562 cells were obtained from^25^. Libraries were sequenced on a MinION device (Oxford Nanopore Technologies) for up to 72 hours.

RNA extracted from mitoribosome-IP samples, from NT or FASTKD5 KD cells, were directly used as input for direct RNA library preparation as described above using 500-700 ng input RNA (without prior poly(A)+ selection).

### Nanopore sequencing data analysis

Live base-calling of nanopore sequencing data was performed with MinKNOW (release 20.10.3 or later). Reads with a base-calling threshold > 7 were converted into DNA sequences by substituting U to T bases prior to alignment. Reads were aligned to the Hela-specific single nucleotide polymorphism-corrected reference hg38 genome (see below) using minimap2^76^ with parameters -ax splice -uf -k14. Multi-mapping reads were included in all downstream analyses. All subsequent analyses were performed using custom python scripts.

For measuring transcript abundance, the alignment BAM file was converted to a BED file using pybedtools^77^ bamtobed with options cigar=True, tag=’NM’. Reads mapping to the mitochondrial genome were extracted using pybedtools intersect with options wo=True, s=True, and a BED file containing the coordinates of all mitochondrial transcripts. Read/transcript pairs that contained the cigar string “N” within 25 nt of the transcript start or end were excluded. Unique reads intersecting each transcript by at least 25 nt were counted. For measuring the abundance of reads mapping to the last 100 nt of the transcript, the same strategy was used with a BED file containing the coordinates of the 3 prime-end most 100 nt for each transcript. For transcripts that are frequently detected in the unprocessed state (e.g. *ATP8-6/COX3*), the abundance of the upstream transcript was corrected to take into account reads that are not long enough to reach it. For each read, the genomic coordinates of the read start and the read end were extracted and intersected with the following genomic features within the transcripts of interest using pybedtools with options wo=True, s=True: the transcript start region (0 to +25 nt transcript start), the transcript end region (-25 to +25 nt from the transcript end), the gene body (+25 nt from transcript start to –25 nt from transcript end) and the 5 prime upstream region (-1500 to 0 nt from transcript start). For the downstream transcript (e.g. *COX3*), the 5 prime-end fraction unprocessed was calculated as the number of reads mapping to this transcript that start in the 5 prime upstream region divided by the number of reads mapping to this transcript that start in the 5 prime upstream region or in the transcript start region. Reads mapping to this transcript that start in the gene body were defined as those lacking information about the 5 prime-end and that potentially extend into the upstream transcript (e.g. *ATP8-6*). The abundance of the upstream transcript was corrected using the following formula: reads mapping to upstream transcript + (fraction unprocessed downstream transcript * number of reads lacking information about 5 prime-end downstream transcripts).

For analysis of processing, the genomic coordinates of the read start and the read end were extracted and intersected with the following genomic features using pybedtools with options wo=True, s=True: the transcript start region (-15 to +50 nt from transcript start), the transcript end region (-15 to +15 nt from transcript end) and the gene body (transcript start to transcript end). Next, the entire reads were intersected with the following genomic features using pybedtools with options wo=True, s=True: the upstream region (-60 to -15 nt from transcript start), the downstream region (+15 to +60 nt from transcript end), the transcript end region (-15 to +15 nt from transcript end) and the gene body (transcript start to transcript end). Reads/feature pairs that overlapped by less than 15 nt or that contained the cigar string “N” within 15 nt of the feature start or end were excluded. Reads that mapped to a protein-coding gene and that ended in the transcript end region of that gene were classified as “processed” at the 3 prime-end. Reads that mapped to a protein-coding gene and to the downstream region of that gene were classified as “unprocessed” at the 3 prime-end. Reads that mapped to a protein-coding gene and that started in the transcript start region of that gene were classified as “processed” at the 5 prime-end. Reads that mapped to a protein-coding gene and to the upstream region of that gene were classified as “unprocessed” at the 5 prime-end.

For identifying the location of reads starts in the NT and *FASTKD5* KD samples, reads that were unprocessed at the 5’-end were extracted using the approach outlined above. The genomic locations of read starts were intersected with the gene body coordinates, as defined above. Read starts locations were categorized based on whether they corresponded to the tRNA or the mRNA gene immediately upstream of the gene of interest (i.e. *tRNA^Leu^* for *ND1*, *tRNA^Met^* for *ND2*, *ATP8-6* for *COX3* and *ND5* for *CYB*), or to genes located further upstream. Split reads, which are likely technical artifacts, were filtered out by converting the BAM file to a BED12 file using bedtools bamtobed and keeping only reads that did not have a comma in the last column.

Poly(A) tail lengths were estimated using nanopolish^41^. For 3 prime-end ligated samples, reads were classified as non-polyadenylated if the qc_tag from nanopolish was “NO_REGION” or polyadenylated if the qc_tag was “PASS” or “ADAPTER”. For poly(A)+, only reads classified as polyadenylated were analyzed. Only reads with the 3 prime-end processing status “processed” were included in poly(A) tail length analyses.

For modeling transcription, the genomic coordinates of the read 3 prime-end were extracted. Using pybedtools with options wo=True, s=True, read ends were classified as “processed” if they intersected with a 25 nt window around the transcript end site or “actively transcribing” if they mapped more than 25 nt upstream of the transcript end site.

### Quantitative gene expression models

To estimate the importance of each of the levels of gene expression that we had acquired kinetic measurements we developed a quantitative gene-expression model (Figure 5) that we defined as follows:

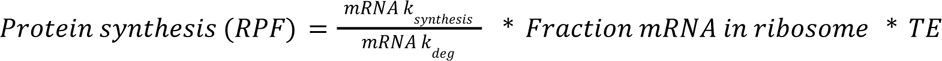

Where the *mRNA k*_*synthesis*_ = 1/(*T*_*initiation*_ + *T*_*elongation*_) from the transcription elongation model defined above, “fraction mRNA in ribosome” as defined above and the 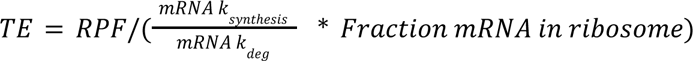. The coefficient of determination from correlating the predicted RPF values from the model to that of measured RPF values was used to measure the predictive power of the model. The model per definition explains all of the variability in RPF. The importance of each gene-expression level was determined by setting the succeeding parameters to 1. E.g. to estimate the predictive value of 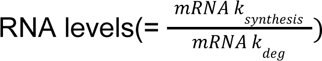 we set both *Fratcion mRAN in ribosome and the TE* to 1.

We also developed a simpler minimal gene-expression model to directly compare nuclear and mitochondrial gene-expression control where 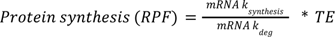 where the *mRAN K*_*synthesis*_ = *RAN levels*/*mRNA k*_*deg*_ and *TE* = *RPF*/*mRNA levels*. RNA-levels were from direct RNA-seq (Figure 1) for mitochondria and TPMs from nuclear-encoded genes. mitoRPF data from HeLa cells were from^3^, HeLa cytoRPF data is from^78^. The predictive power was determined as the coefficient of determination from correlating the predicted RPF values from the model to that of measured RPF values, either from the same HeLa data set (Model) or a HeLa replicate (Replicate). The importance of each gene-expression node was again determined by sequentially setting the TE and then *mRAN K*_synthesis_ parameters to 1. All rates, RPF, TPM, and direct RNA-seq values can be found in the supplemental tables. For the nuclear side we excluded two outliers, ATP5MC3 and ATP5F1E, out of 75 genes total as they single-handedly depressed the RPF correlation.

---

**Supplemental Figure 1.**
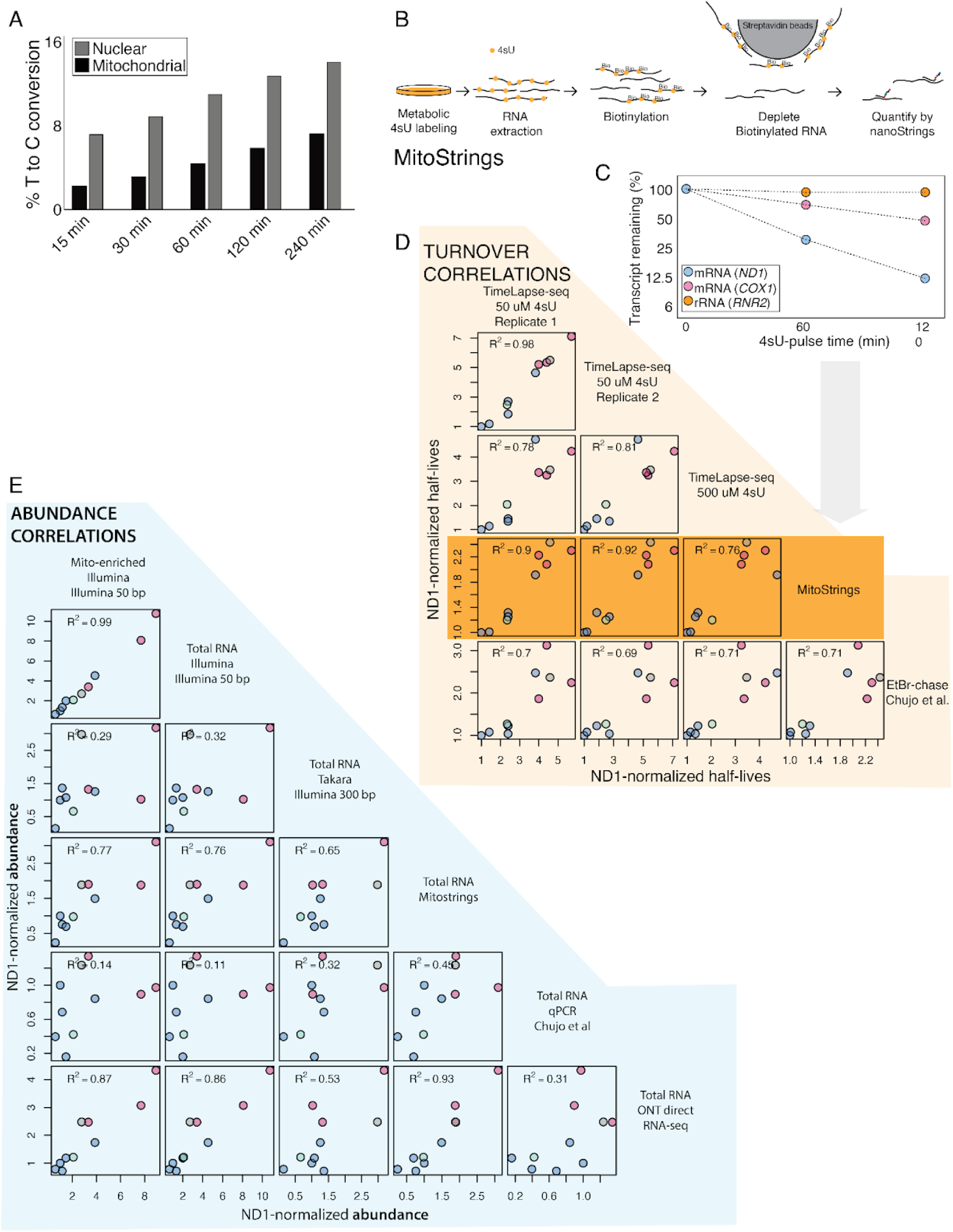
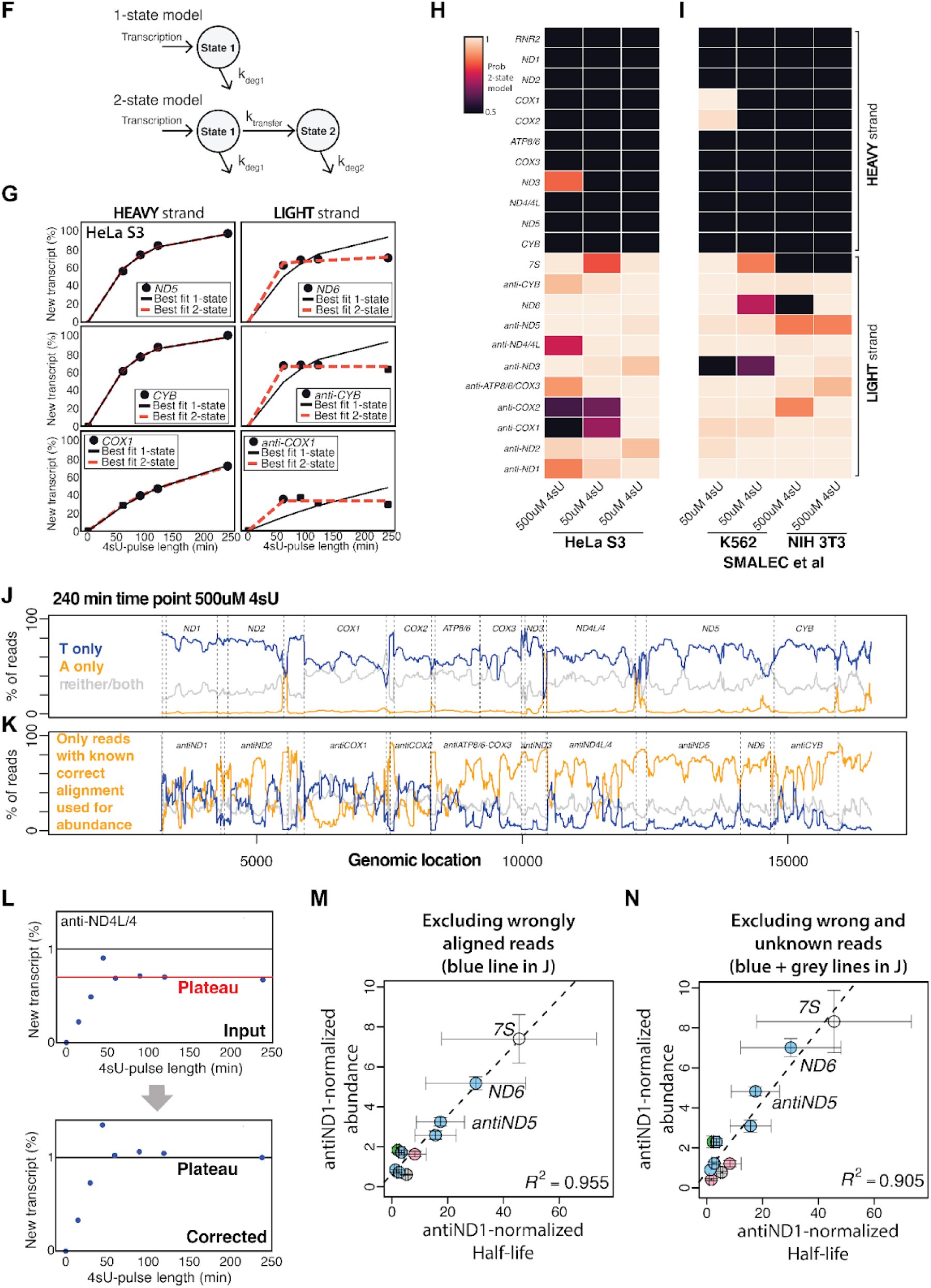
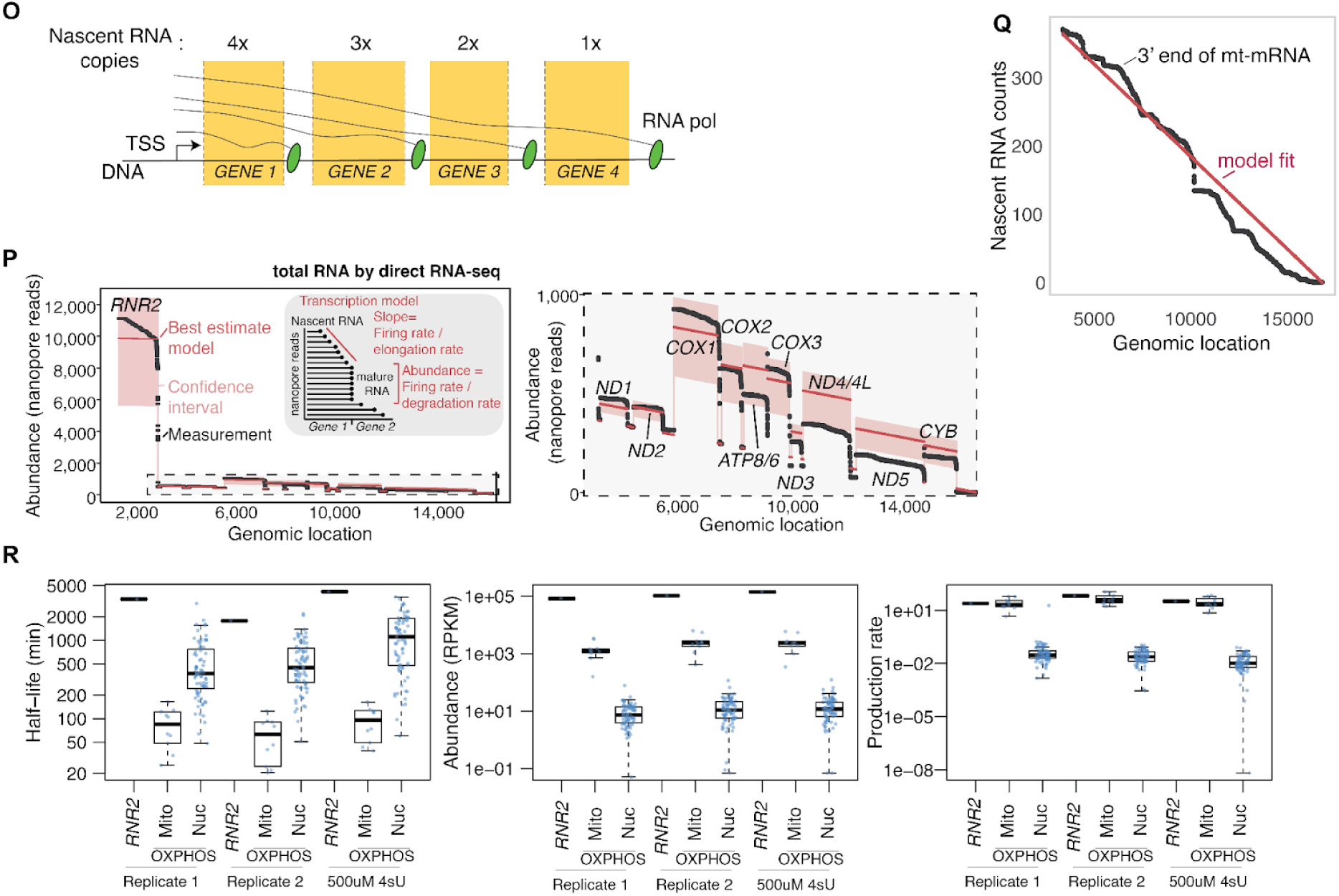
A) Barplot showing the fraction T’s that have been converted to C’s by Time-Lapse chemistry and reverse transcription after labeling HeLa cells with 500uM 4sU. Nuclear-encoded transcripts consistently have a higher conversion rate than mitochondrial-encoded transcripts at the same time point. B) 4sU labeled RNA is biotinylated and then removed after incubation with streptavidin beads leaving only pre-existing “old” RNA. The decrease in native/unlabeled RNA over time can be measured by the MitoStrings assay (see material and methods). C) Representative example profiles from one of three biological replicates. D) Correlation between turnover measurement using MitoStrings (from C), with TimeLapse-seq using 50 or 500uM 4sU or previously published data from HeLa cells deploying ethidium bromide (EtBr) chase experiments with qPCR as a readout^18^. E) Correlation between abundance estimates using different readout methods. F) We deployed a 2-state Markov model to detect RNA with turnover profiles that significantly differed from a simple exponential model. G) Example profiles from light and heavy strand transcripts. Light strand transcript can be seen to plateau before reaching maximum turnover. The light strand-encoded transcriptome consists of one mRNA, *ND6*, the 7S regulatory RNA, eight tRNAs, and numerous antisense (or ‘mirror’) RNAs with unknown functions. H) Systematic analysis of the occurrence of a plateau shows that this behavior is consistent but limited to the light-strand encoded transcripts. I) The same behavior can be seen in previously published data sets with RNA from different cell lines deploying different indexing and sequencing chemistries but the same library prep kit^25^. J and K) The plateau coincides with a significant amount of reads aligning to the wrong strand as judged by the 4sU-induced mismatches showing up at the wrong strand. The heavy strand transcriptome J) displays very few incorrectly aligned reads (A to G mismatches - orange line) while the light strand K) shows significant amounts of misaligned reads (T to C mismatches - blue line). We reasoned these wrongly aligned reads are caused by strand-switching or index hopping during RNA-seq library generation^79^ and estimate that ∼3% of the heavy-strand RNAs are converted into contaminating reads mapping to the light strand. The high abundance of the heavy strand and low abundance of light strand encoded transcripts makes this a directional problem mainly impacting the light strand-encoded transcripts. L) Light strand turnover profiles can be corrected by normalizing each profile so that the last time point is set to 100% new. This moves the plateau up to where we would expect it to be in case the plateau reflects a fully turned-over transcript (bottom panel). The difference between the top and bottom panels is assumed to be caused by wrongly aligned reads that cannot contain the expected mismatches caused by 4sU. M and N) Using corrected turnover profiles (as in L) to calculate half-lives as well as corrected abundance estimates based on all reads aligning to the light strand minus the known wrongly aligned reads (M, blue line in K) or known and unknown (N, we cannot determine if reads lacking a 4sU induced mismatch (gray line in K) are aligned to the correct strand or not). Shown is the average of *antiND1* normalized values and error bars show standard deviations. O) The nascent (actively transcribed) transcriptome has a TSS-proximal gene bias because, at any one point in time, more RNA polymerases will have traversed proximal genes than distal genes. As the mitochondrial transcriptome is rapidly turned over a significant fraction of all mt-mRNA are nascent and thus this bias could impact overall RNA levels. P) Zoom in on the mRNA highlighted in the dashed line box in the first panel (same as Figure 1K). Q) Transcription elongation rates were estimated based on the slope of 3’ ends of mt-mRNA that are actively transcribed. The black line shows reads from nanopore direct RNA sequencing that are considered nascent RNA as their 3’ ends align to the gene bodies as opposed to the 3’ cleavage sites across the heavy strand. The red line shows the model fit estimating transcription elongation rate (see material and methods). R) Same as in Figure 1 D-F) but showing all three experiments and individual data points. All data in the figure are from HeLa cells except those in I) which include data from NIH-3T3 and K562 cells.

**Supplemental Figure 2.**
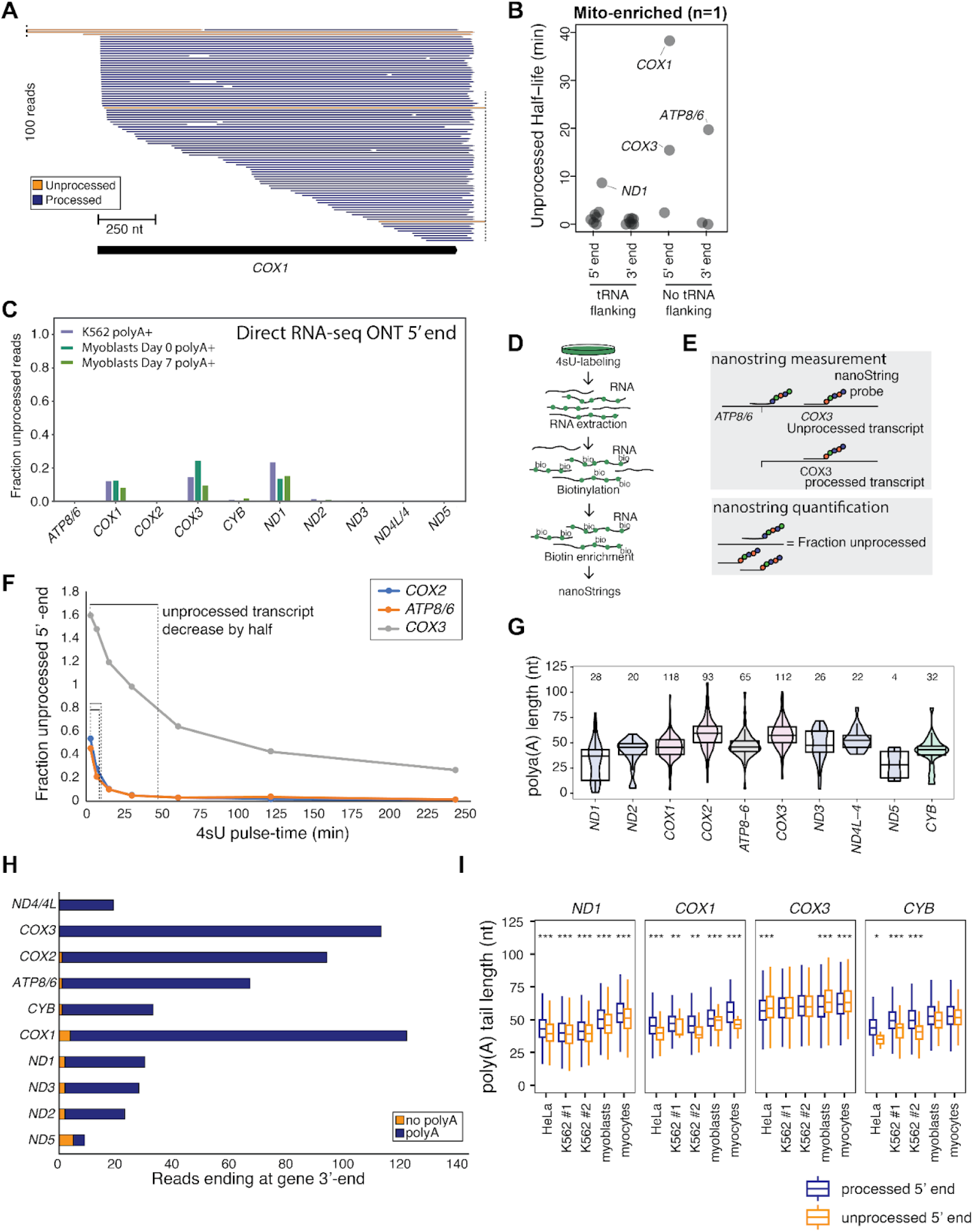
A) 100 randomly sampled reads from nanopore direct RNA-seq that align to *COX1*. Orange reads are unprocessed at the 5’ and/or the 3’ end, and blue reads are processed or undetermined in cases where the read does not reach the end of the transcript. B) The amount of time required until half of the newly synthesized transcripts have been processed at a specific site using nanopore direct RNA-seq data from a mitochondria-enriched sample. Processing sites are subdivided into 3’ or 5’ ends of the mRNA as well as into groups of canonical (tRNA-flanking) or non-canonical. C) Bar plot showing the fraction of unprocessed 5’ ends in poly(A)-selected RNA sequenced by nanopore direct RNA-seq. Data is from K562 cells and human myoblasts before or after 7 days of differentiation towards myotubes. D) Cartoon showing the experimental procedure for the data shown in E). E) MitoStrings can be used to estimate the fraction of unprocessed transcript by applying probes that bind either gene internally or overlap a processing site. F) Data for three processing junctions using the experimental design as shown in D) and analyzed as in E). G) Poly(A) tail length from nanopore direct RNA-seq experiment from total rRNA depleted RNA. H) All reads that end at the 3’ end of a gene, from the same experiment as in G), and whether they have been polyadenylated or not. I) Poly(A) tail length as a function of 5’ end processing status for four cell lines. All data in the figure are from HeLa cells if not stated otherwise.

**Supplemental Figure 3.**
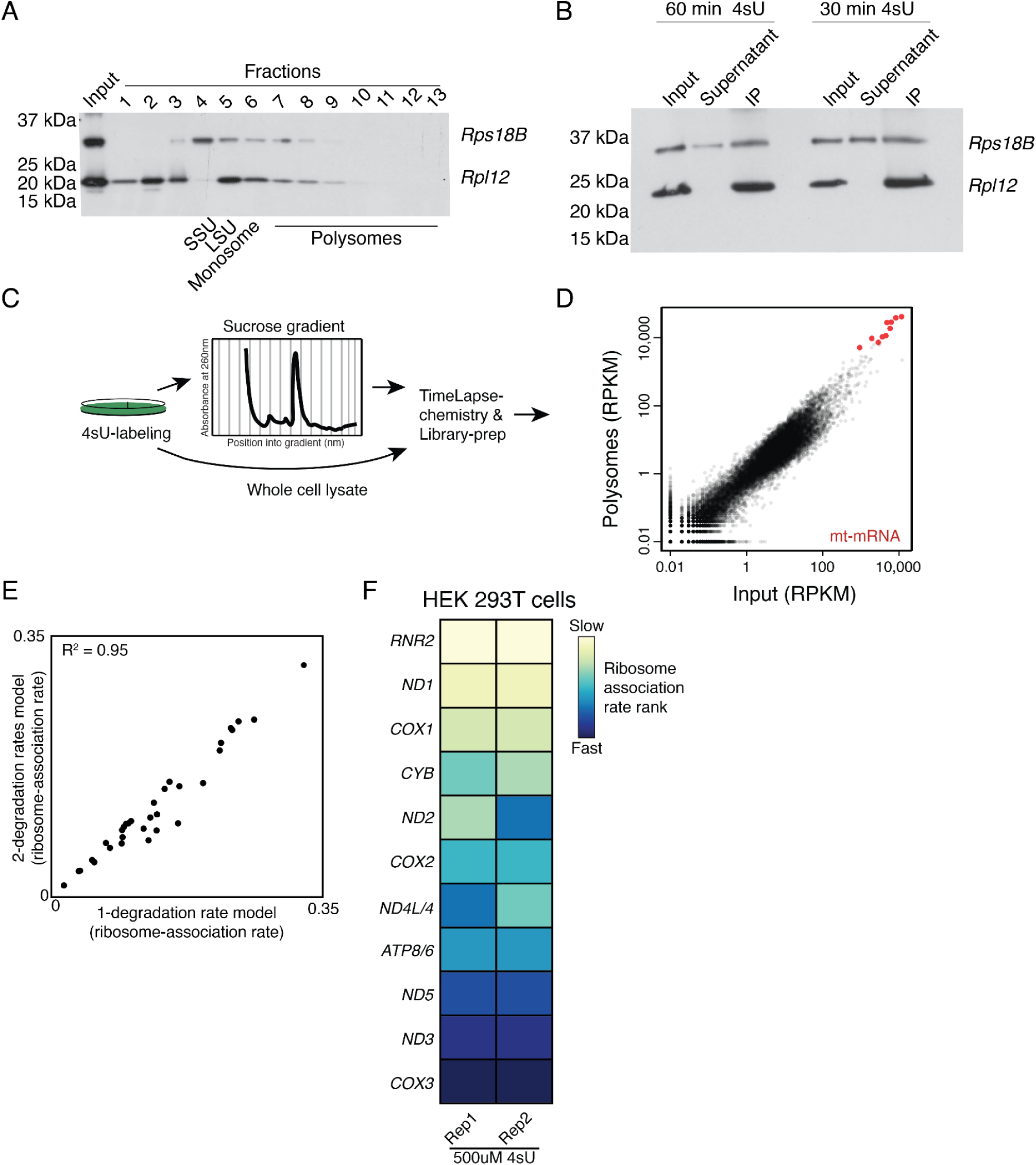
A) Western blot of two ribosomal proteins, one small subunit (*Rps18B*) and one large subunit (Rpl12), from proteins extracted out of a sucrose gradient such as in C). The fractions where the majority of the small ribosomal subunit (SSU), large ribosomal subunit (LSU), monosome and polysome reside have been annotated. B) Western Blot for the same ribosomal proteins in A) but here showing the result of immunoprecipitation of the ribosome out of pooled gradient fractions from two 4sU labeling time points. All of Rpl12 (the bait) from the input comes down in the IP fraction and none remains in the supernatant. For Rps18B the majority comes down in the IP while a fraction remains in the supernatant. C) Cartoon of the experimental design using only sucrose gradient to enrich for mitochondrial ribosome-associated RNA. D) Results from the experiment outlined in C). RPKMs from libraries made from RNA in the input or extracted from pooled sucrose gradient fractions (fractions 6-13) show that there is little enrichment for mt-mRNA or mt-rRNA overall cellular mRNA or antisense (non-translated) mt-transcripts. E) Scatter plot comparing the ribosome-association rates (from all three replicates) using either the simpler mathematical model assuming the same degradation rate for both the ribosome-associated RNA and the unbound RNA (1-degradation rate model) or a slightly more complex model allowing for independent degradation rates for the two states (2-degradation rate model, see material and methods). Both models result in similar estimates of the ribosome association rates. F) Heatmap showing the transcripts ranked by their ribosome-association half-lives in each replicate using HEK293T cells. All data in the figure are from HeLa cells if not stated otherwise.

**Supplemental Figure 4.**
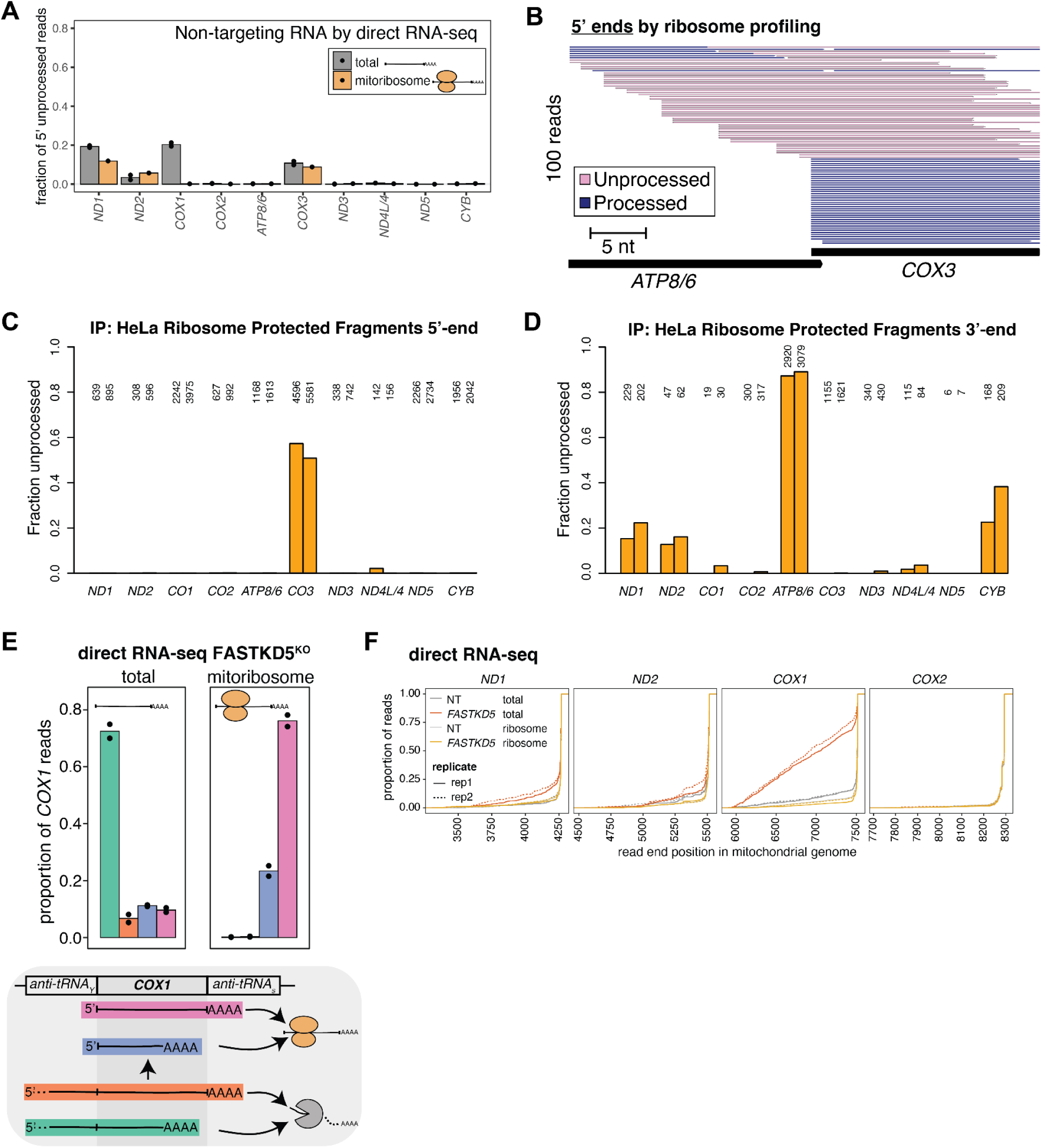
A) Comparing the fraction of 5’-end unprocessed mRNA in total and ribosome-associated mRNA in the HEK293T cells. Each dot or triangle represents data from one of two biological replicates. Triangles and darker-colored bars indicate the fraction of reads that end on the tRNA closest to the 5’ end of the analyzed mRNA B) Subsampling of 100 ribosome-protected fragments (RPF) with 5’ ends within -30 and +3 nt from the 5’ end of *COX3*. The scale bar shows the distance in nucleotides, and the color code refers to the processing status of each read. C) Quantification of all ribosome-protected fragments that align to the 5 prime ends of mitochondrial genes from immunoprecipitated ribosomes^3^. The plot shows the fraction of the reads that are unprocessed (i.e. are orange in B)) over all reads aligning to the 5 prime end of the gene (I.e. blue and orange reads in B)). D) Same as in B) but for 3 prime ends. When compared to the non-immunoprecipitated RPF from HeLa cells in Figure 4, there are no major differences. E) The proportion of *COX1* mRNA in *FASTKD5* KD cells that is processed at either the 5’ or 3’-end as judged by direct RNA-seq of total or ribosome-associated RNA. The scheme below summarizes the finding that only 5’ end processed mRNA associate with the ribosome. F) 3’-end locations of reads from direct RNA-sequencing experiments using either total or ribosome-associated RNA as input. Only *COX1* mRNA in the FASTKD5 KD shows a large fraction of gene-internal 3’-ends. Panels A), E), and F) show data from HEK293T cells, panels B), C), and D) show data from HeLa cells.

## References

1. Isaac, R.S., McShane, E., and Churchman, L.S. (2018). The Multiple Levels of Mitonuclear Coregulation. Annu. Rev. Genet. 52, 511–533.

2. Couvillion, M.T., Soto, I.C., Shipkovenska, G., and Churchman, L.S. (2016). Synchronized mitochondrial and cytosolic translation programs. Nature 533, 499–503.

3. Soto, I., Couvillion, M., Hansen, K.G., McShane, E., Moran, J.C., Barrientos, A., and Churchman, L.S. (2022). Balanced mitochondrial and cytosolic translatomes underlie the biogenesis of human respiratory complexes. Genome Biol. 23, 170.

4. Köhler, F., Müller-Rischart, A.K., Conradt, B., and Rolland, S.G. (2015). The loss of LRPPRC function induces the mitochondrial unfolded protein response. Aging 7, 701–717.

5. Suomalainen, A., and Battersby, B.J. (2017). Mitochondrial diseases: the contribution of organelle stress responses to pathology. Nat. Rev. Mol. Cell Biol. 10.1038/nrm.2017.66.

6. Aloni, Y., and Attardi, G. (1971). Symmetrical in vivo transcription of mitochondrial DNA in HeLa cells. Proc. Natl. Acad. Sci. U. S. A. 68, 1757–1761.

7. Ojala, D., Montoya, J., and Attardi, G. (1981). tRNA punctuation model of RNA processing in human mitochondria. Nature 290, 470.

8. Ojala, D., Merkel, C., Gelfand, R., and Attardi, G. (1980). The tRNA genes punctuate the reading of genetic information in human mitochondrial DNA. Cell 22, 393–403.

9. Rackham, O., and Filipovska, A. (2022). Organization and expression of the mammalian mitochondrial genome. Nat. Rev. Genet. 10.1038/s41576-022-00480-x.

10. Tan, B.G., Mutti, C.D., Shi, Y., Xie, X., Zhu, X., Silva-Pinheiro, P., Menger, K.E., Díaz-Maldonado, H., Wei, W., Nicholls, T.J., et al. (2022). The human mitochondrial genome contains a second light strand promoter. Mol. Cell 82, 3646–3660.e9.

11. Bullerwell, C.E., and Gray, M.W. (2004). Evolution of the mitochondrial genome: protist connections to animals, fungi and plants. Curr. Opin. Microbiol. 7, 528–534.

12. Formaggioni, A., Luchetti, A., and Plazzi, F. (2021). Mitochondrial Genomic Landscape: A Portrait of the Mitochondrial Genome 40 Years after the First Complete Sequence. Life 11. 10.3390/life11070663.

13. Foury, F., Roganti, T., Lecrenier, N., and Purnelle, B. (1998). The complete sequence of the mitochondrial genome of Saccharomyces cerevisiae. FEBS Lett. 440, 325–331.

14. Li, G.-W., Burkhardt, D., Gross, C., and Weissman, J.S. (2014). Quantifying absolute protein synthesis rates reveals principles underlying allocation of cellular resources. Cell 157, 624–635.

15. Sasarman, F., Brunel-Guitton, C., Antonicka, H., Wai, T., Shoubridge, E.A., and LSFC Consortium (2010). LRPPRC and SLIRP interact in a ribonucleoprotein complex that regulates posttranscriptional gene expression in mitochondria. Mol. Biol. Cell 21, 1315–1323.

16. Gelfand, R., and Attardi, G. (1981). Synthesis and turnover of mitochondrial ribonucleic acid in HeLa cells: the mature ribosomal and messenger ribonucleic acid species are metabolically unstable. Mol. Cell. Biol. 1, 497–511.

17. Mercer, T.R., Neph, S., Dinger, M.E., Crawford, J., Smith, M.A., Shearwood, A.-M.J., Haugen, E., Bracken, C.P., Rackham, O., Stamatoyannopoulos, J.A., et al. (2011). The human mitochondrial transcriptome. Cell 146, 645–658.

18. Chujo, T., Ohira, T., Sakaguchi, Y., Goshima, N., Nomura, N., Nagao, A., and Suzuki, T. (2012). LRPPRC/SLIRP suppresses PNPase-mediated mRNA decay and promotes polyadenylation in human mitochondria. Nucleic Acids Res. 40, 8033–8047.

19. Piechota, J., Tomecki, R., Gewartowski, K., Szczesny, R., Dmochowska, A., Kudła, M., Dybczyńska, L., Stepien, P.P., and Bartnik, E. (2006). Differential stability of mitochondrial mRNA in HeLa cells. Acta Biochim. Pol. 53, 157–168.

20. Herzog, V.A., Reichholf, B., Neumann, T., Rescheneder, P., Bhat, P., Burkard, T.R., Wlotzka, W., von Haeseler, A., Zuber, J., and Ameres, S.L. (2017). Thiol-linked alkylation of RNA to assess expression dynamics. Nat. Methods 14, 1198–1204.

21. Schofield, J.A., Duffy, E.E., Kiefer, L., Sullivan, M.C., and Simon, M.D. (2018). TimeLapse-seq: adding a temporal dimension to RNA sequencing through nucleoside recoding. Nat. Methods 15, 221–225.

22. Riml, C., Amort, T., Rieder, D., Gasser, C., Lusser, A., and Micura, R. (2017). Osmium-Mediated Transformation of 4-Thiouridine to Cytidine as Key To Study RNA Dynamics by Sequencing. Angew. Chem. Int. Ed Engl. 56, 13479–13483.

23. Burger, K., Mühl, B., Kellner, M., Rohrmoser, M., Gruber-Eber, A., Windhager, L., Friedel, C.C., Dölken, L., and Eick, D. (2013). 4-thiouridine inhibits rRNA synthesis and causes a nucleolar stress response. RNA Biol. 10, 1623–1630.

24. Jürges, C., Dölken, L., and Erhard, F. (2018). Dissecting newly transcribed and old RNA using GRAND-SLAM. Bioinformatics 34, i218–i226.

25. Smalec, B.M., Ietswaart, R., Choquet, K., McShane, E., West, E.R., and Stirling Churchman, L. (2022). Genome-wide quantification of RNA flow across subcellular compartments reveals determinants of the mammalian transcript life cycle. bioRxiv, 2022.08.21.504696. 10.1101/2022.08.21.504696.

26. Wolf, A.R., and Mootha, V.K. (2014). Functional genomic analysis of human mitochondrial RNA processing. Cell Rep. 7, 918–931.

27. Siira, S.J., Spåhr, H., Shearwood, A.-M.J., Ruzzenente, B., Larsson, N.-G., Rackham, O., and Filipovska, A. (2017). LRPPRC-mediated folding of the mitochondrial transcriptome. Nat. Commun. 8, 1532.

28. Wu, Z., Ietswaart, R., Liu, F., Yang, H., Howard, M., and Dean, C. (2016). Quantitative regulation of FLC via coordinated transcriptional initiation and elongation. Proc. Natl. Acad. Sci. U. S. A. 113, 218–223.

29. Ietswaart, R., Rosa, S., Wu, Z., Dean, C., and Howard, M. (2017). Cell-Size-Dependent Transcription of FLC and Its Antisense Long Non-coding RNA COOLAIR Explain Cell-to-Cell Expression Variation. Cell Syst 4, 622–635.e9.

30. Ruzzenente, B., Metodiev, M.D., Wredenberg, A., Bratic, A., Park, C.B., Cámara, Y., Milenkovic, D., Zickermann, V., Wibom, R., Hultenby, K., et al. (2012). LRPPRC is necessary for polyadenylation and coordination of translation of mitochondrial mRNAs. EMBO J. 31, 443–456.

31. Singh, V., Itoh, Y., Huynen, M.A., and Amunts, A. (2022). Activation mechanism of mitochondrial translation by LRPPRC-SLIRP. bioRxiv, 2022.06.20.496763. 10.1101/2022.06.20.496763.

32. Sultana, S., Solotchi, M., Ramachandran, A., and Patel, S.S. (2017). Transcriptional fidelities of human mitochondrial POLRMT, yeast mitochondrial Rpo41, and phage T7 single-subunit RNA polymerases. J. Biol. Chem. 292, 18145–18160.

33. Yu, H., Xue, C., Long, M., Jia, H., Xue, G., Du, S., Coello, Y., and Ishibashi, T. (2018). TEFM Enhances Transcription Elongation by Modifying mtRNAP Pausing Dynamics. Biophys. J. 115, 2295–2300.

34. Posse, V., Shahzad, S., Falkenberg, M., Hällberg, B.M., and Gustafsson, C.M. (2015). TEFM is a potent stimulator of mitochondrial transcription elongation in vitro. Nucleic Acids Res. 43, 2615–2624.

35. Wuarin, J., and Schibler, U. (1994). Physical isolation of nascent RNA chains transcribed by RNA polymerase II: evidence for cotranscriptional splicing. Mol. Cell. Biol. 14, 7219–7225.

36. Pandya-Jones, A., and Black, D.L. (2009). Co-transcriptional splicing of constitutive and alternative exons. RNA 15, 1896–1908.

37. Wachutka, L., Caizzi, L., Gagneur, J., and Cramer, P. (2019). Global donor and acceptor splicing site kinetics in human cells. Elife 8. 10.7554/eLife.45056.

38. Wan, Y., Anastasakis, D.G., Rodriguez, J., Palangat, M., Gudla, P., Zaki, G., Tandon, M., Pegoraro, G., Chow, C.C., Hafner, M., et al. (2021). Dynamic imaging of nascent RNA reveals general principles of transcription dynamics and stochastic splice site selection. Cell 184, 2878–2895.e20.

39. Temperley, R.J., Wydro, M., Lightowlers, R.N., and Chrzanowska-Lightowlers, Z.M. (2010). Human mitochondrial mRNAs--like members of all families, similar but different. Biochim. Biophys. Acta 1797, 1081–1085.

40. Rackham, O., Mercer, T.R., and Filipovska, A. (2012). The human mitochondrial transcriptome and the RNA-binding proteins that regulate its expression. Wiley Interdiscip. Rev. RNA 3, 675–695.

41. Workman, R.E., Tang, A.D., Tang, P.S., Jain, M., Tyson, J.R., Razaghi, R., Zuzarte, P.C., Gilpatrick, T., Payne, A., Quick, J., et al. (2019). Nanopore native RNA sequencing of a human poly(A) transcriptome. Nat. Methods 16, 1297–1305.

42. Bratic, A., Clemente, P., Calvo-Garrido, J., Maffezzini, C., Felser, A., Wibom, R., Wedell, A., Freyer, C., and Wredenberg, A. (2016). Mitochondrial Polyadenylation Is a One-Step Process Required for mRNA Integrity and tRNA Maturation. PLoS Genet. 12, e1006028.

43. Ng, K.Y., Lutfullahoglu Bal, G., Richter, U., Safronov, O., Paulin, L., Dunn, C.D., Paavilainen, V.O., Richer, J., Newman, W.G., Taylor, R.W., et al. (2022). Nonstop mRNAs generate a ground state of mitochondrial gene expression noise. Sci Adv 8, eabq5234.

44. Nicholson, A.L., and Pasquinelli, A.E. (2019). Tales of Detailed Poly(A) Tails. Trends Cell Biol. 29, 191–200.

45. Remes, C., Khawaja, A., Pearce, S.F., Dinan, A.M., Gopalakrishna, S., Cipullo, M., Kyriakidis, V., Zhang, J., Dopico, X.C., Yukhnovets, O., et al. (2023). Translation initiation of leaderless and polycistronic transcripts in mammalian mitochondria. Nucleic Acids Res. 10.1093/nar/gkac1233.

46. Bogenhagen, D.F., Ostermeyer-Fay, A.G., Haley, J.D., and Garcia-Diaz, M. (2018). Kinetics and Mechanism of Mammalian Mitochondrial Ribosome Assembly. Cell Rep. 22, 1935–1944.

47. Montoya, J., Ojala, D., and Attardi, G. (1981). Distinctive features of the 5’-terminal sequences of the human mitochondrial mRNAs. Nature 290, 465–470.

48. Lagouge, M., Mourier, A., Lee, H.J., Spåhr, H., Wai, T., Kukat, C., Silva Ramos, E., Motori, E., Busch, J.D., Siira, S., et al. (2015). SLIRP Regulates the Rate of Mitochondrial Protein Synthesis and Protects LRPPRC from Degradation. PLoS Genet. 11, e1005423.

49. Christian, B.E., and Spremulli, L.L. (2010). Preferential Selection of the 5′-Terminal Start Codon on Leaderless mRNAs by Mammalian Mitochondrial Ribosomes. J. Biol. Chem. 285, 28379–28386.

50. Antonicka, H., and Shoubridge, E.A. (2015). Mitochondrial RNA Granules Are Centers for Posttranscriptional RNA Processing and Ribosome Biogenesis. Cell Rep. 10.1016/j.celrep.2015.01.030.

51. Ohkubo, A., Van Haute, L., Rudler, D.L., Stentenbach, M., Steiner, F.A., Rackham, O., Minczuk, M., Filipovska, A., and Martinou, J.-C. (2021). The FASTK family proteins fine-tune mitochondrial RNA processing. PLoS Genet. 17, e1009873.

52. Martin, M., Cho, J., Cesare, A.J., Griffith, J.D., and Attardi, G. (2005). Termination factor-mediated DNA loop between termination and initiation sites drives mitochondrial rRNA synthesis. Cell 123, 1227–1240.

53. Morgenstern, M., Peikert, C.D., Lübbert, P., Suppanz, I., Klemm, C., Alka, O., Steiert, C., Naumenko, N., Schendzielorz, A., Melchionda, L., et al. (2021). Quantitative high-confidence human mitochondrial proteome and its dynamics in cellular context. Cell Metab. 10.1016/j.cmet.2021.11.001.

54. Bekker-Jensen, D.B., Kelstrup, C.D., Batth, T.S., Larsen, S.C., Haldrup, C., Bramsen, J.B., Sørensen, K.D., Høyer, S., Ørntoft, T.F., Andersen, C.L., et al. (2017). An Optimized Shotgun Strategy for the Rapid Generation of Comprehensive Human Proteomes. Cell Syst 4, 587–599.e4.

55. Morisaki, T., and Stasevich, T.J. (2018). Quantifying Single mRNA Translation Kinetics in Living Cells. Cold Spring Harb. Perspect. Biol. 10. 10.1101/cshperspect.a032078.

56. Hausser, J., Mayo, A., Keren, L., and Alon, U. (2019). Central dogma rates and the trade-off between precision and economy in gene expression. Nat. Commun. 10, 68.

57. Siwiak, M., and Zielenkiewicz, P. (2013). Transimulation - protein biosynthesis web service. PLoS One 8, e73943.

58. Richter-Dennerlein, R., Oeljeklaus, S., Lorenzi, I., Ronsör, C., Bareth, B., Schendzielorz, A.B., Wang, C., Warscheid, B., Rehling, P., and Dennerlein, S. (2016). Mitochondrial Protein Synthesis Adapts to Influx of Nuclear-Encoded Protein. Cell 167, 471–483.e10.

59. Fuchs, G., Voichek, Y., Benjamin, S., Gilad, S., Amit, I., and Oren, M. (2014). 4sUDRB-seq: measuring genomewide transcriptional elongation rates and initiation frequencies within cells. Genome Biol. 15, R69.

60. Shah, P., Ding, Y., Niemczyk, M., Kudla, G., and Plotkin, J.B. (2013). Rate-limiting steps in yeast protein translation. Cell 153, 1589–1601.

61. Farge, G., and Falkenberg, M. (2019). Organization of DNA in Mammalian Mitochondria. Int. J. Mol. Sci. 20. 10.3390/ijms20112770.

62. Shpilka, T., and Haynes, C.M. (2018). The mitochondrial UPR: mechanisms, physiological functions and implications in ageing. Nat. Rev. Mol. Cell Biol. 19, 109–120.

63. Khalimonchuk, O., Bird, A., and Winge, D.R. (2007). Evidence for a pro-oxidant intermediate in the assembly of cytochrome oxidase. J. Biol. Chem. 282, 17442–17449.

64. Song, J., Herrmann, J.M., and Becker, T. (2021). Quality control of the mitochondrial proteome. Nat. Rev. Mol. Cell Biol. 22, 54–70.

65. Lane, N., and Martin, W. (2010). The energetics of genome complexity. Nature 467, 929–934.

66. Lane, N. (2014). Bioenergetic constraints on the evolution of complex life. Cold Spring Harb. Perspect. Biol. 6, a015982.

67. Gomes, A.P., Price, N.L., Ling, A.J.Y., Moslehi, J.J., Montgomery, M.K., Rajman, L., White, J.P., Teodoro, J.S., Wrann, C.D., Hubbard, B.P., et al. (2013). Declining NAD(+) induces a pseudohypoxic state disrupting nuclear-mitochondrial communication during aging. Cell 155, 1624–1638.

68. Houtkooper, R.H., Mouchiroud, L., Ryu, D., Moullan, N., Katsyuba, E., Knott, G., Williams, R.W., and Auwerx, J. (2013). Mitonuclear protein imbalance as a conserved longevity mechanism. Nature 497, 451–457.

69. López-Otín, C., Blasco, M.A., Partridge, L., Serrano, M., and Kroemer, G. (2022). Hallmarks of aging: An expanding universe. Cell. 10.1016/j.cell.2022.11.001.

70. Hanahan, D., and Weinberg, R.A. (2011). Hallmarks of Cancer: The Next Generation. Cell 144, 646–674.

71. Akaike, H. (1974). A new look at the statistical model identification. IEEE Trans. Automat. Contr. 19, 716–723.

72. Sin, C., Chiarugi, D., and Valleriani, A. (2016). Degradation Parameters from Pulse-Chase Experiments. PLoS One 11, e0155028.

73. Stefan Isaac, R., Tullius, T.W., Hansen, K.G., Dubocanin, D., Couvillion, M., Stergachis, A.B., and Stirling Churchman, L. (2022). Single-nucleoid architecture reveals heterogeneous packaging of mitochondrial DNA. bioRxiv, 2022.09.25.509398. 10.1101/2022.09.25.509398.

74. Drexler, H.L., Choquet, K., Merens, H.E., Tang, P.S., Simpson, J.T., and Churchman, L.S. (2021). Revealing nascent RNA processing dynamics with nano-COP. Nat. Protoc. 16, 1343–1375.

75. Mayer, A., and Churchman, L.S. (2016). Genome-wide profiling of RNA polymerase transcription at nucleotide resolution in human cells with native elongating transcript sequencing. Nat. Protoc. 11, 813–833.

76. Li, H. (2018). Minimap2: pairwise alignment for nucleotide sequences. Bioinformatics 34, 3094–3100.

77. Dale, R.K., Pedersen, B.S., and Quinlan, A.R. (2011). Pybedtools: a flexible Python library for manipulating genomic datasets and annotations. Bioinformatics 27, 3423–3424.

78. Wu, C.C.-C., Zinshteyn, B., Wehner, K.A., and Green, R. (2019). High-Resolution Ribosome Profiling Defines Discrete Ribosome Elongation States and Translational Regulation during Cellular Stress. Mol. Cell 73, 959–970.e5.

79. Costello, M., Fleharty, M., Abreu, J., Farjoun, Y., Ferriera, S., Holmes, L., Granger, B., Green, L., Howd, T., Mason, T., et al. (2018). Characterization and remediation of sample index swaps by non-redundant dual indexing on massively parallel sequencing platforms. BMC Genomics 19, 332.

